# Heparan sulfate is essential for Drosophila FGF export

**DOI:** 10.64898/2026.03.24.714045

**Authors:** Guilherme Oliveira Barbosa, Christian Solis-Calero, Thomas B. Kornberg

## Abstract

Binding of Fibroblast growth factor (FGF) to a heparan sulfate proteoglycan (HSPG) is required for paracrine FGF signaling. To improve our understanding of FGF:HSPG association, we developed a method to monitor export of the Drosophila FGF ortholog Branchless (Bnl) *in vivo*. We detected Bnl on the surface of approximately 10% of Bnl-producing cells, but Bnl on the surface of cells depleted of HS was much reduced. HS depletion also non-autonomously decreased the activity of cytonemes that extend from cells that receive Bnl. These results are consistent with the idea that Bnl export to the cell surface is regulated, that intracellular binding of an HSPG to Bnl in producing cells is essential for export, and that cells that take up Bnl actively participate in its release from producing cells.

**Summary:** Levels of FGF exported to the surface of FGF-expressing cells are dependent on intracellular heparan sulfate proteoglycans.

## Introduction

Fibroblast Growth Factors (FGFs) are signaling proteins in metazoans which have important roles in development and tissue organization (Itoh and Ornitz, 2004). Vertebrates have 22 FGFs; *Drosophila melanogaster* has three - Branchless (Bnl), Pyramus, and Thisbe. The structural hallmark of FGFs is an approximately 120 amino acid β-trefoil domain composed of 12 β-strands. Although the N- and C-terminal regions that flank the conserved FGF domain are varied, fifteen of the FGFs have a heparin sulfate binding site (HBS) (Goetz and Mohammadi, 2013). These HBS sites do not share a defined sequence, but all have closely spaced lysine and arginine residues and a positive electrostatic potential that promotes ionic interaction between heparan sulfate (HS) and FGF (Xu and Esko, 2014). The HBS contributes to the formation of a tertiary complex composed of FGF, FGF receptor (FGFR) and a HS proteoglycan (HSPG) (Robinson et al., 2005; Schlessinger et al., 2000). The FGFs in the fifteen-member, HBS-containing subfamily are the “canonical FGFs”; they are all thought to be exported from the cells that produce them.

HS polymers are covalently linked to proteoglycans; they are synthesized by exostosin enzymes and are modified by sulfotransferases and epimerases. The affinity of HS for the positive electrical potential of an FGF HBS is dependent on two factors: the negative charge of HS sulfated saccharides (N-acetyl-glycosaminoglycan / Glucuronic acid) and polysaccharide chain flexibility which is enhanced by epimerases that convert glucuronic acid residues to iduronic acid (Xu and Esko, 2014).

FGF signal transduction is mediated by tyrosine kinase FGF receptors. Vertebrates have four; Drosophila has two - Breathless (Btl) and Heartless (Htl). Binding by FGF together with HS induces FGFR dimerization and triggers intracellular signaling through various pathways, including Ras-MAPK, PI3K-AKT, and PLCγ1 (Goetz and Mohammadi, 2013). In vertebrates, these pathways control developmental outcomes such as neurogenesis and epithelial branching (i.e.: lung, kidney, mammary gland, salivary gland and prostate) (Bellusci et al., 1997; Huang et al., 2005; Makarenkova et al., 2009; Qiao et al., 2001; Steinberg et al., 2005; Sumbal and Koledova, 2019). In Drosophila, Btl FGFR mediates Bnl signaling in tracheal branching morphogenesis (Sato and Kornberg, 2002; Sutherland et al., 1996), while the Htl FGFR is required for the development of the heart and several other mesodermal tissues (Beiman et al., 1996; Patel et al., 2022).

During development of the Drosophila embryo, invaginations of the ectoderm form tracheal pits, and Bnl expressed by clusters of nearby epidermal and mesenchymal cells activates Btl-initiated signal transduction in tracheal pit cells to induce the formation of the tracheal system (Sutherland et al., 1996). Later during the third larval instar (L3), Bnl produced by the wing disc promotes the development of the Air Sac Primordium (ASP), an epithelial tube which extends from a disc-associated tracheal branch (Sato and Kornberg, 2002). Extensive investigations have established that wing disc-produced Bnl moves to the ASP via Btl-bearing cytonemes that extend from the ASP and contact Bnl-producing wing disc cells (Du et al., 2018). These signaling-specialized filopodia are called cytonemes (Ramírez-Weber and Kornberg, 1999).

Cytonemes project from both the Bnl source cells of the disc and from Bnl receiving cells in the ASP, providing both specificity and directionality to the FGF paracrine signaling (Du et al., 2018). Activation of Bnl signaling in the ASP also induces a positive feedback response that enhances reception of Bnl, whereby Bnl-receiving ASP cells extend more Btl-containing cytonemes (Du et al., 2018). Cytonemes extend over as many as 12-15 cell diameters from the distal tip of the ASP, consistent with the range of FGF paracrine signaling in flies and vertebrates (Shimokawa et al., 2011; Venero Galanternik et al., 2015).

Similarities between cell-cell signaling at the tips of neuronal axons and signaling at cytoneme contacts are more than superficial: shared functions features include proteins that constitute vertebrate glutamatergic synapses (e.g., the glutamate receptor, glutamate transporter, voltage-gated calcium channel, synaptotagmin, synaptobrevin) and functionalities that include ionic current transsynaptic activation (Huang et al., 2019)., The extension of Btl-containing cytonemes depends on the expression of the

Drosophila glypican Dally-like protein (Dlp) in the wing disc (Huang and Kornberg, 2016). HSPGs are also needed for stabilization of neuronal dendrites (Poe et al., 2017). The underlying mechanisms involved in the HS-dependence of contact-mediated paracrine signaling by neurons and non-neuronal cells remains unknown.

To investigate the role of HS in cytoneme-mediated signaling, we exploited features of Drosophila ASP development that make it possible to independently manipulate gene expression in the ASP and wing disc and to observe the affected processes in real time. We found that in wing discs depleted of HS, Bnl signaling in the ASP was reduced and ASP cytonemes were not normal. Although ASP cytonemes extended and retracted over HS-depleted regions of the wing disc, their dynamics were abnormal. We also developed an *in vivo* method to assess export of Bnl to the cell surface, and observed that <10% of cells that express Bnl in the wing disc had detectable levels of cell surface Bnl, that cells depleted of HS had reduced levels of cell surface Bnl, and that the levels of a mutant Bnl with an altered HBS were reduced on the surface of cells with normal HS production. These results confirm the importance of HS-HSB affinity in Bnl signaling and reveal an essential intracellular function for HSPG in Bnl export. We suggest that these functions of HSPGs may not be unique to the FGFs but may also be important for other signaling proteins,

## Results

### A HS role in Bnl paracrine signaling

We genetically depleted HS production in the wing disc to investigate the role of HSPGs in Bnl signaling from the disc to the ASP. To address the fact that different HSPGs may have partially redundant and compensatory activities that complicate interpretation of HSPG mutant phenotypes, we focused on HS production because all HSPGs are dependent on the same enzymes of the HS biosynthetic pathway and are all affected by reduced HS functionality. We first measured expression levels of the nine HS biosynthetic pathway genes by qPCR analysis of RNA isolated from wing discs. The transcript levels varied over an approximately 10-fold range; *Sulfated* (*sulf1*) RNA was the most abundant, at ∼1.6% the amount of *Act5C* RNA (Fig. 1A). We next analyzed the genes by RNAi knockdown in the dorsal compartment of the wing disc (*ap*>GAL4), and assessed ASP morphology by measuring its length and width (Fig. 1B). Knockdown of genes involved in 3-O sulfation (*hs3sta, hs3stb*) had minimal effect on ASP shape.

**Figure 1:**
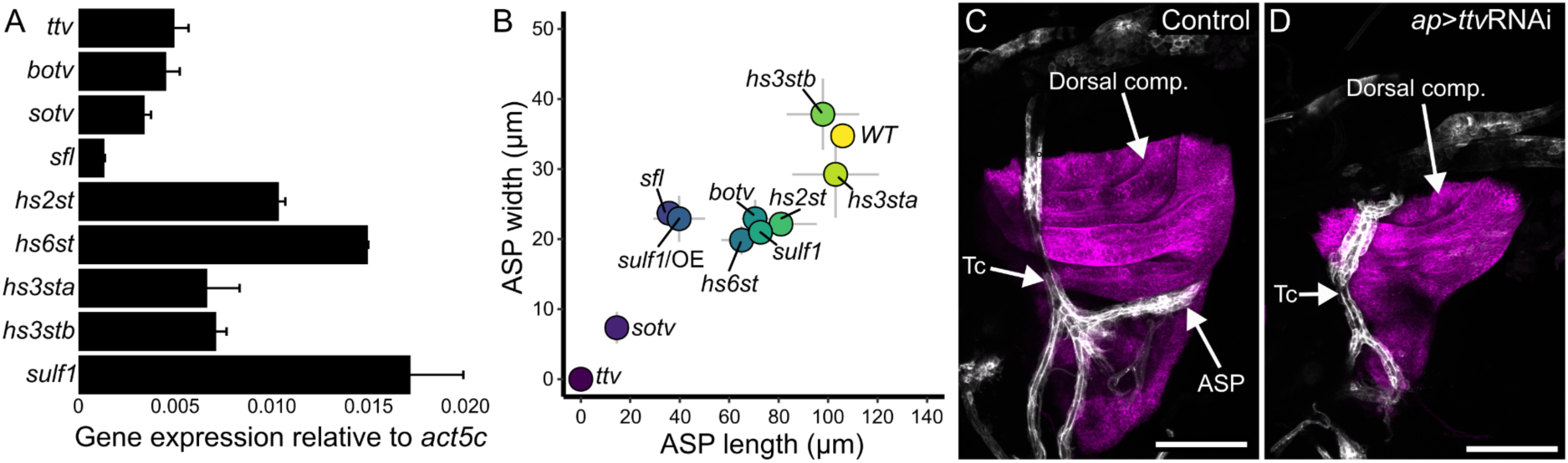
ASP morphogenesis requires HS synthesis in wing disc cells. (A) Third instar wing disc expresses enzymes of HS synthesis (*ttv*, *botv*, *sotv*) and modification (*sfl*, *hs2st*, *hs6st*, *hs3sta*, *hs3stb*, *sulf1*). Values: average from two experimental pools of 15 wing discs; bars: expression relative to Act5c; error bars: standard deviation. (B) Plot of ASP (marked with *btl>*CD4:GFP fluorescence) width and length in genotypes with *ap>*RNAi knockdown of *ttv*, *botv*, *sotv*, *sfl*, *hs2st*, *hs6st*, *hs3sta*, *hs3stb*, *sulf1*, and over-expression of sulfatese1 (*ap>*SULF1). Dot locations: value means (n=5(*ttv)*, 6(*botv)*, 16(*sotv)*, 2*(sfl)*, 4*(hs2st)*, 6*(hs6st)*, 4*(hs3sta)*, 5*(hs3stb)*, 10*(sulf1)*, 8*(SULF1)*; gray bars: standard deviation (width – y axis, length – x axis). (C,D) ASP (white) develops juxtaposed to the wing disc dorsal compartment (magenta) in control (*btl>*CD4:GFP *ap>*CD4:mIFP) but not in *ttv* knockdown (*btl>*CD4:GFP *ap>*CD4:mIFP *ap>*ttvRNAi) L3 animals. Scale bars 100 µm.

Knockdown of the genes in involved 2-O (*hs2st*) and 6-O (*hs6st*) sulfation affected both ASP length and width, and similar effects were observed after knockdown of *bother-of-ttv* (*botv*), which is involved in HS synthesis, or *sulf1*, which is responsible for 6-O sulfation removal. Kamimura et al (2006) reported evidence for *hs2st* and *hs6st* redundancy, consistent with the idea than knockout of either gene does not generate a null phenotype. RNAi expression directed against *sulfateless* (*sfl*), which is involved in N-sulfation, and *sulf1* overexpression (SULF1) had greater effects on ASP growth.

Knockdown of either *sister-of-ttv* (*sotv*) or *tout-velu* (*ttv*), which encode two enzymes that initiate HS synthesis, had the most extreme effects on the ASP (Fig. 1C,D). These results are consistent with our previous work which showed that Dlp produced in the wing disc is required by the ASP (Huang and Kornberg, 2016). The results are not in agreement with the Yan and Lin, 2007 study (Yan and Lin, 2007) which concluded that the ASP is not affected by *sfl* or *dally/dlp* mutant clones in the wing disc. In our opinion, the ASP morphologies in the published images are not normal.

To determine if HS produced in the wing disc is required to activate the ERK/MAPK pathway in the ASP, we analyzed ASP development and used antibody directed against phosphorylated ERK to monitor dpERK in ASP preparations from wild types and from animals in which *ttv*RNAi was expressed in the wing disc. We examined ASPs in late L3 animals following 24, 48, or 72 hours of *ap>*ttvRNAi expression, and observed reductions in ASP growth (Fig. 2A-I), cytoneme number (Fig. 2J), and cytoneme length (Fig. 2K) that increased in severity with the longer periods of expression. In controls and in animals after 24 hours of *ttv*RNAi expression, dpERK was present in cells at the ASP tip closest to the basal surface of the wing disc, but was absent after 48 hours of expression (Fig. 2L,L’,M,M’). The photomicrographs in Figures 1C,D and 2A-D show that *ap*>GAL4 induction of *ttv* RNAi reduced the size of both the ASP and wing disc.

**Figure 2:**
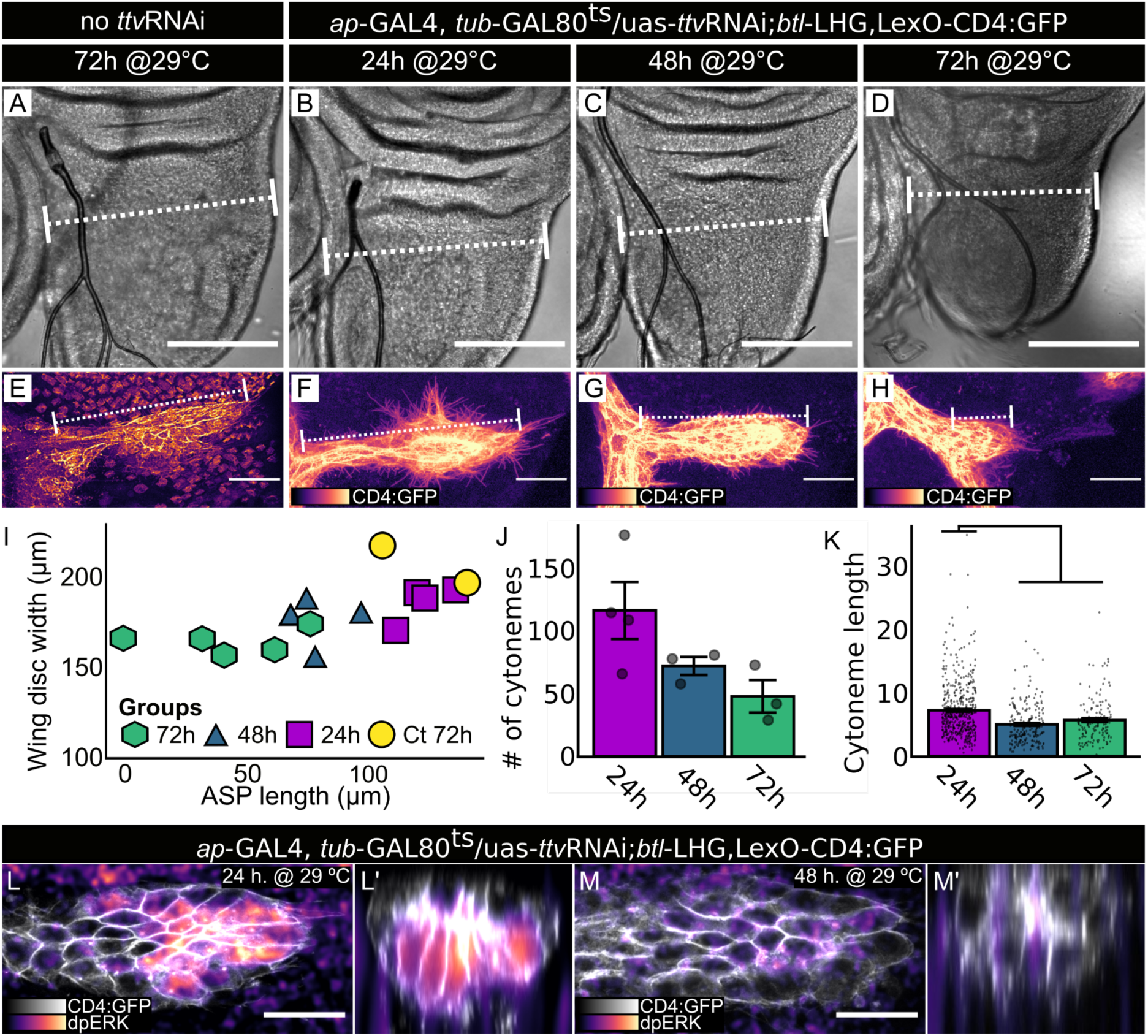
*ttv* RNAi expression in the wing disc reduces ASP growth, cytoneme numbers and length. (A-K) *ttv*RNAi expression in wing disc dorsal compartment (Gal80^ts^, *ap>*ttvRNAi) in control (A,E (n=2)) and experimentals (B-D, F-H) for indicated times prior to dissection at L3: (B) 24hr (n=4); (C), 48hr (n=4); (D) 72hr (n=5). (A-D) DIC images; dotted lines indicate site of measurement for disc width; scale bar, 30 µm. (E-H) Images of ASP (*btl>*CD4:GFP) at indicated times of *ttv*RNAi expression prior to dissection at L3; scale bar, 100 µm. (I) Plot of average wing disc width and ASP length for (A-I) preparations; length differences between 48 and 72 hour preparations of control and experimentals is statistically significant (ANOVA: Df=3, F=8.075. Tukey’s HDS: Ct-48h *P*=0.03; Ct-72h *P*=0.003). (J) Numbers of cytonemes per ASP after 24 (n=4), 48 (n=3), and 72 (n=3) hrs of *ap*>ttvRNAi expression (ANOVA: Df=2, F=3.946. Tukey’s HDS: no significant difference). (K) Lengths of cytonemes quantified in (J) (ANOVA: Df=3, F=25.39. Tukey’s HDS: 24h-48h *P<*0.001; 24h-72h *P<*0.001). (L) Normal dpERK staining in ASP tip (white, frontal view) after 24 hr *ttv* RNAi expression (magenta scale; Gal80^ts^, *btl>*CD4:GFP, *ap>*ttvRNAi); (L’) transverse projection. (M,M’) Reduced dpERK staining after 48 hr *ttv*RNAi expression (images as in (L,L’)). Scale bars (L-M), 20 µm.

The absence of dpERK in ASPs analyzed after 48 hours of *ttv*RNAi expression is consistent with the idea that HS-modified HSPGs in the wing disc are essential for Bnl signaling in the ASP. These findings are also consistent both with previous work showing that lack of HS sulfation in the wing disc and ASP causes defects in ASP morphogenesis (Kamimura et al., 2006), and with our earlier work which concluded that Bnl signal transduction in the ASP requires the HSPG Dlp in wing disc cells (Huang and Kornberg, 2016).

Because Bnl signaling is not required for wing disc development (Sato and Kornberg, 2002), the stunted growth of wing discs that express *ap>*ttvRNAi (Fig.1D and Fig. 2D) is likely due to deficits in signaling systems other than Bnl - such as Dpp and Hh. To characterize the effects of ttvRNAi expression further, we expressed ttvRNAi in the dorsal compartment (*ap*>ttvRNAi; Fig. 3A,D,E,H,I). We monitored Dpp signaling with *dad*-GFP, a transcriptional reporter of Dpp signal transduction (Fig. 3B-E), and monitored Hh signaling by immunohistochemical staining for Ptc (Fig. 3G-I), which is upregulated by Hh signal transduction. Both the fluorescence of *dad*-GFP and α-Ptc immunostaining were reduced in discs that expressed ttvRNAi in the dorsal compartment.

**Figure 3:**
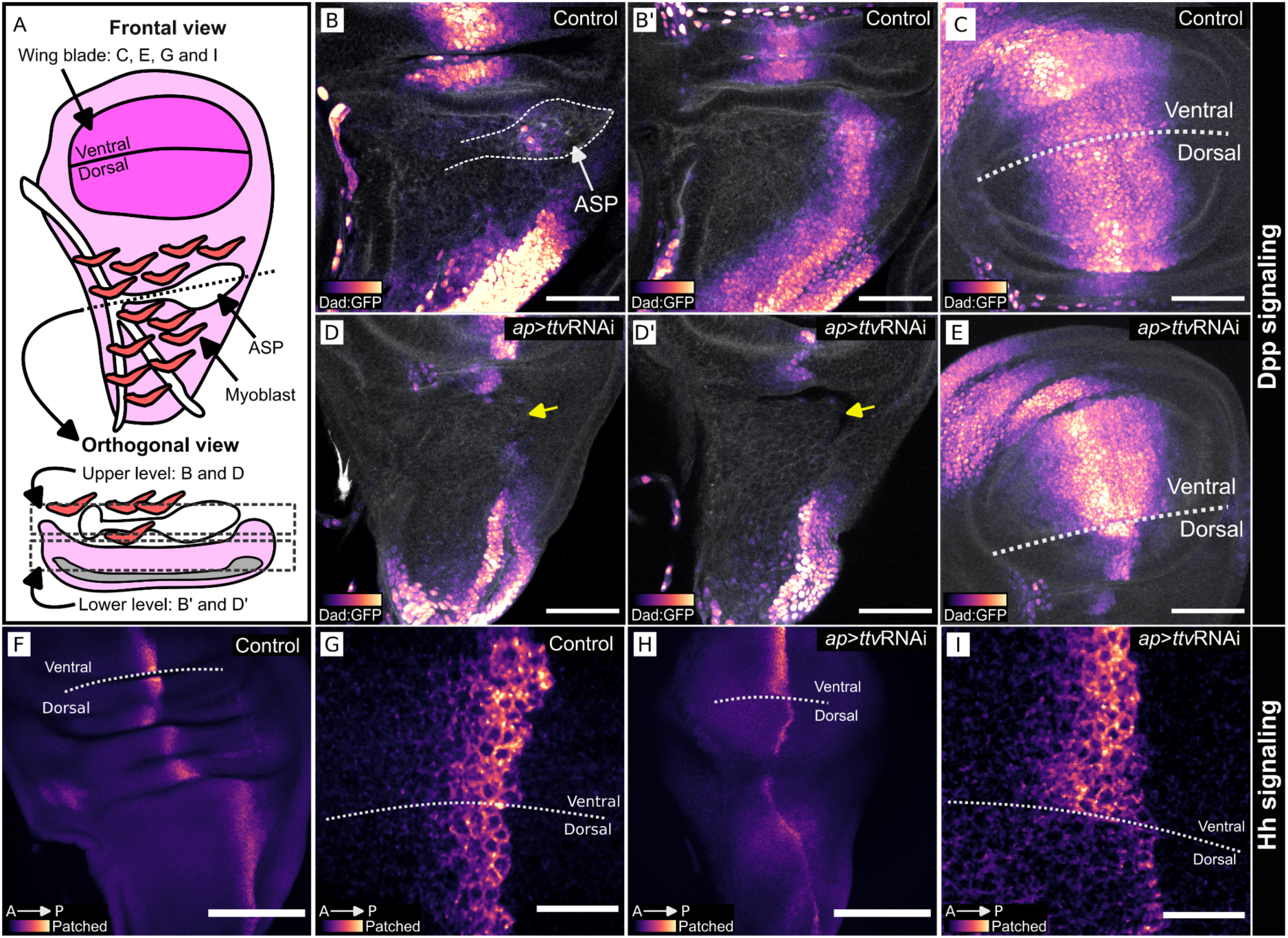
*ttv* knockdown in wing disc dorsal compartment reduces Dpp and Hh signaling. (A) Wing disc schematic (frontal and orthogonal views) with myoblasts (red) and ASP (white). (B-E) L3 discs imaged for Dad:GFP (magenta scale), reporter for Dpp signaling, and Phalloidin (white), stained with Alexa Flour^TM^ 647. (B-C) Dad:GFP in wing disc and ASP; (B) upper and (B’) lower orthogonal optical sections of dorsal compartment, (C) frontal view of wing blade primordium. (D-E) Dad:GFP fluorescence is reduced in the dorsal compartment of the wing disc after *ttv* RNAi expression (*ap>*ttvRNAi), shown in (D) upper and (D’) lower orthogonal optical sections of dorsal compartment and (E) frontal view of the wing blade primordium. (D,D’) yellow arrow indicates the normal location of ASP). (F-I) Staining for Hh signaling reporter Ptc of control wing disc (F,G), showing enhanced staining at A/P compartment border. (H-I) Wing discs expressing *ttv*RNAi in dorsal compartment (*ap>*ttvRNAi) have reduced Ptc dorsal compartment staining. Scale bars: (B-E) 50 µm. (F,H) 100 µm, (G,I) 20 µm.

We expressed ttvRNAi in either the entire anterior compartment (*ci*>ttvRNAi; Fig. 4A), in anterior compartment cells at the compartment border (*ptc*>ttvRNAi; Fig. 4B), or in the entire posterior compartment (*hh*>ttvRNAi; Fig. 4C). Whereas the proximal/distal lengths of the anterior and posterior compartments in wild type wing discs are similar, expression of ttvRNAi in the anterior compartment (either *ci*>ttvRNAi or *ptc*>ttvRNAi) reduced the size of the anterior compartment and almost eliminated ASP development, but did not affect the posterior compartment in a comparable way. Expression of ttvRNAi in the posterior compartment (*hh*>ttvRNAi) did not reduce the size of the anterior compartment markedly, but it reduced the size of the posterior compartment and limited ASP growth such that the ASP extended up to but not beyond the anteroposterior border (Fig. 4D,D’). These results are consistent with the idea that HSPGs produced in the wing disc are required for Dpp and Hh signaling for the growth of both anterior and posterior wing disc compartments. They do not indicate whether the effects on the ASP implicate Dpp and Hh signaling deficits in the ASP.

**Figure 4:**
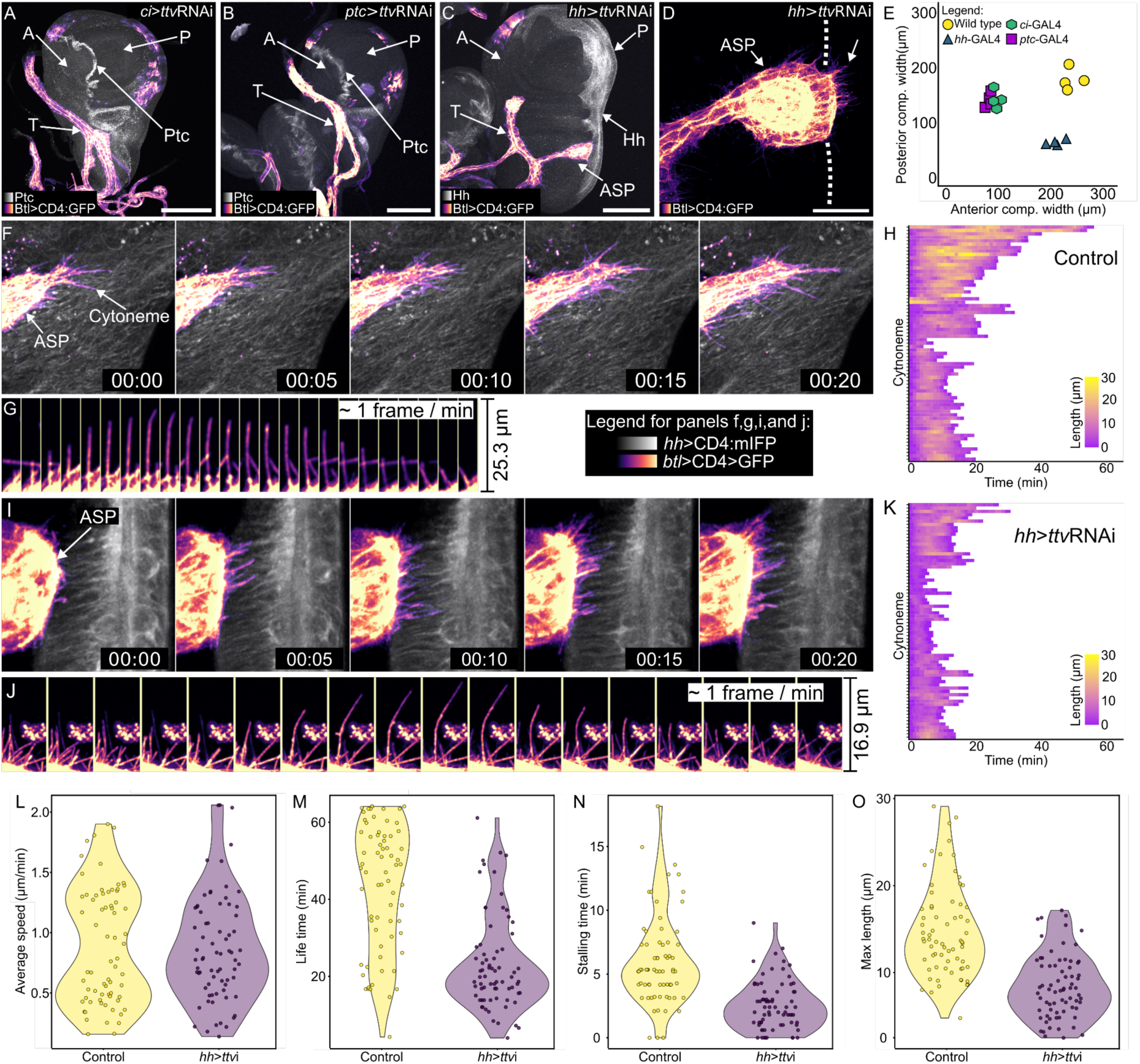
Life time, stalling time and maximum length of ASP are reduced in areas reduced HS. (A-D) *ttv* knockdown in: (A) A compartment (*ci*-Gal4, α-Ptc antibody staining (white) locates A/P compartment border); (B) A cells at A/P compartment border (*ptc*-Gal4, UAS-CD4:mIFP (white)); (C,D) P compartment (*hh*-Gal4, UAS-Cd4:mIFP (white)). *ttv* knockdown reduced disc compartment size (E) and either blocked (A,B) or truncated (C) ASP growth. (T – tracheal branch) Scale bars: (A-C) 100 µm, (D) 30 µm. (F-K) Time lapse recordings (1 hour, ∼1 frame/min) of ASP cytonemes extending over wild type wing disc P compartment (F,G,H) or P compartment of disc expressing *ttv* RNAi (I,J,K). (H,K) Lengths of individual cytonemes (n=66 (H), 63 (K); colored scale) extending and retracting for indicated time (x-axis). (L) Average speed (*t*-test, one-tail, *P*=0.423) of cytonemes extending and retracting over wild type wing disc cells and *ttv* expressing cells are similar. (M) Lifetime, (N) stalling time, and (O) maximum length of cytonemes was reduced over disc cells expressing *ttv* RNAi (*t*-test, one-tail, *P*<0.001).

### Essential roles of HS and HSPGs in wing disc and ASP development

The known Drosophila HSPGs are Division abnormally delayed (Dally), Dally-like protein (Dlp), Syndecan (Sdc), Neurexin IV (NrxIV), Carrier of Wingless (Cow) (Chang and Sun, 2014; Zhang et al., 2018), and Perlecan (Pcan/Trol) (Park et al., 2003). With the exception of Pcan, these HSPGs are expressed by the wing disc, and except for Cow and Pcan, which are secreted to the ECM, they are membrane-bound. Because inhibition of HS synthesis in the wing disc by *ttv* knockdown (*ap*>ttvRNAi) is likely to affect the wing disc-produced HSPGs, the severe effects we observed for ASP morphogenesis (Figs. 1,2) are most likely caused by the absence of HS chains on some or all of these HSPGs and by the essential functions of the wing disc HSPGs for Bnl signaling to the ASP.

The Pcan HSPG is a constituent of the Drosophila ECM. Most Pcan associated with the wing disc is produced by the fat body and/or by hemocytes (Zang et al., 2015) (Fig. 5), and is taken up from the hemolymph. We reduced HS synthesis in the fat body (*lsp2*>ttvRNAi) and hemocytes (*hml*>ttvRNAi) to determine whether the HS-modified proteins produced by the fat body and hemocytes are important for ASP morphogenesis (Fig. 5A). We found that whereas ASP morphogenesis was normal when HS synthesis was reduced in hemocytes (Fig. 5B,D), the ASP was abnormal when HS was depleted in the fat body (Fig. 5C,D). These results are consistent with the idea that Pcan from the hemolymph is essential for interactions between the ASP and wing disc but does not distinguish whether the requisite Pcan is associated with the ECM or is directly associated with the disc cells.

**Figure 5:**
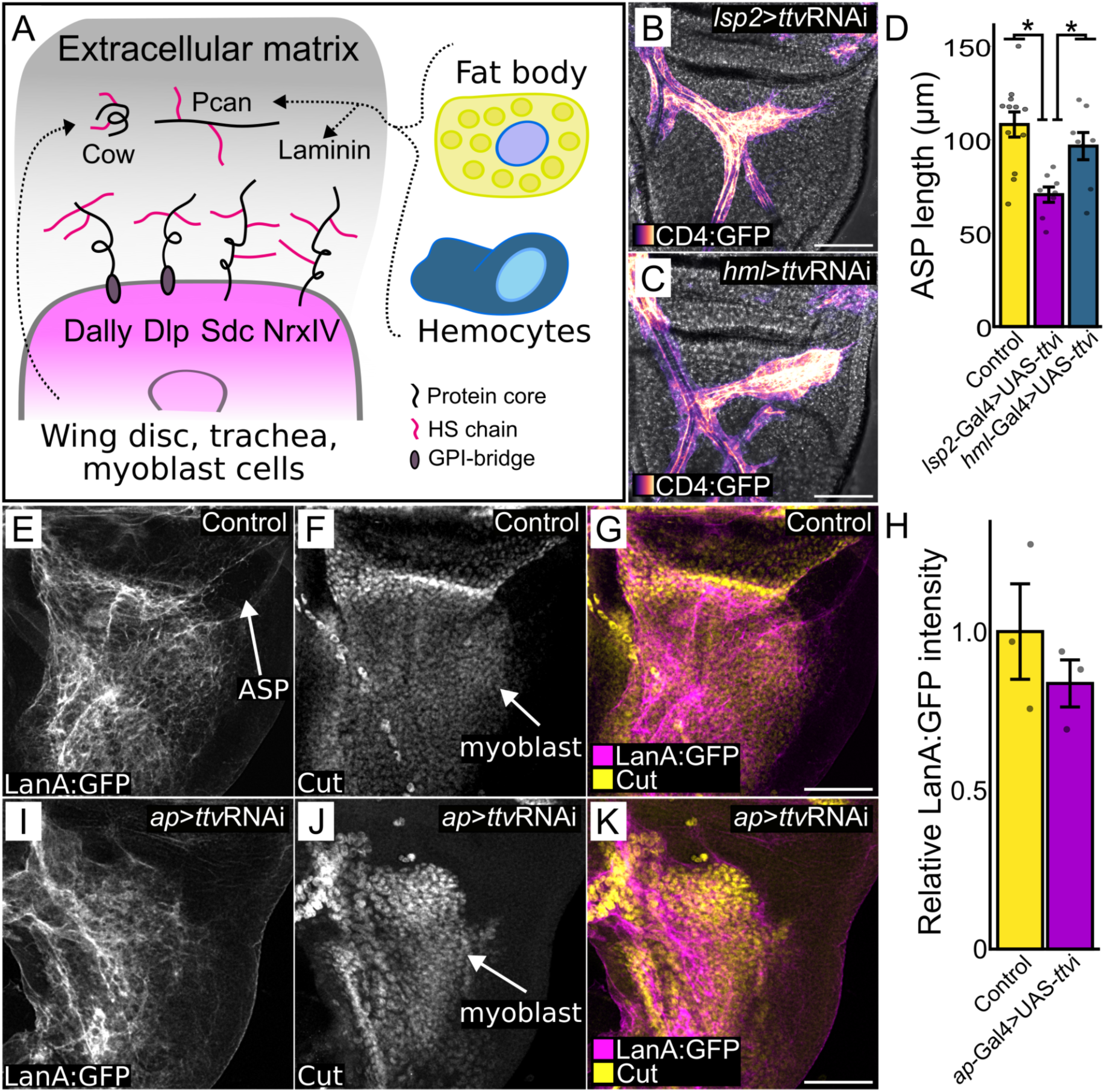
Fat body-produced HS required for ASP and disc-produced HS required for myoblasts. (A) Cartoon depicting wing disc and ECM and HSPGs produced by the disc or received from fat body and hemocytes. (B,C) ASP (magenta scale) in phenotypes with *ttv* knockdown in the fat body (stunted growth; *lsp2*-ttvRNAi) (B) or hemocytes (normal growth; *hml*-ttvRNAi) (C). (D) ASP length in genotypes with either *ttv* knockdown) in fat body (normal) or hemocytes (reduced) (ANOVA: Df=2 F=8.6594. Tukey’s HDS: Control-Lsp2 *P<*0.001; Lsp2-Hml *P<*0.05). (E-K) Laminin associated with wing disc dorsal compartment ECM (Lan:GFP fluorescence, gray scale). (F,J) Reduced number of myoblasts (anti-Cut staining, gray scale) in *ttv* knockdown discs (J) relative to control (F). (H) Average Lan:GFP fluorescence intensity (n=3; dots represent individual values) relative to control in control and ttvRNAi discs (*t*-test, one-tail, no significance). Scale bars: (B,C,E-J) 100 µm.

We probed the integrity of the wing disc ECM under conditions of HS depletion by monitoring Laminin, another ECM constituent that is produced by the fat body and is taken up from the hemolymph (Pastor-Pareja and Xu, 2011). We did not find that *ttv* knockdown of HS in the dorsal wing disc (*ap*>ttvRNAi) affected the presence or distribution of Laminin (Fig. 5E-K). We interpret this result as consistent with the idea that the gross structure of the ECM was not affected by lack of wing disc HS.

### ASP cytoneme dynamics, HS and Bnl signaling

Several studies have described cytoneme extension and retraction dynamics (Bischoff et al., 2013; Chen et al., 2017; Du et al., 2018; González-Méndez et al., 2017; Patel et al., 2022; Simon et al., 2021). Du et al., 2018 reports that cytonemes that extend between the wing disc and ASP had a lifetime of approximately 10-20 minutes, with growth rates of approximately 1.1 μ/minute (Du et al., 2018). We investigated whether cytoneme dynamics were affected by the absence of HSPGs in cells the ASP cytonemes contact – whether the failure of the ASP to extend over P compartment cells that lack HSPGs (*hh*>ttvRNAi) might be related to impaired functionality of ASP cytonemes.

Bnl is produced in the disc by a discrete group of approximately 100 cells that are mostly in the anterior compartment (Du et al., 2018; Sato and Kornberg, 2002) (Supplemental Fig. 1). In discs without P compartment HS-modified HSPGs (*hh*>ttvRNAi), the ASP, which normally extends posteriorly across the A/P compartment border, extended over the A compartment Bnl-expressing cells, but not over the P compartment cells (Fig. 4C,D). Images of cytonemes at the ASP tip revealed that although the ASP did not cross the A/P compartment border (Fig. 4D), ASP cytonemes did cross the border to reach P compartment cells. We developed a method to observe and quantify the dynamics of these cytonemes and to compare their behaviors to cytonemes in control discs (Barbosa and Kornberg, 2022).

We observed ASP cytonemes for 60 minutes at 1 frame/minute and analyzed the cytoneme lifetime, length, rate of extension and retraction, and stalling frequency (length change < 50% in two consecutive time points,). The average speed of extension and retraction of the cytonemes was ∼0.9 µm/min in both experimental and control groups (Fig. 4F-L), comparable to the rate reported in (Du et al., 2018). However, in the HSPG-defective discs, the average lifetime of the cytonemes decreased by almost one half, from ∼44 min to ∼23 min, the average stalling time decreased from 6.3 min to 2.9 min, and the average maximum length decreased from 15.1 µm to 8.8 µm (Fig. 4M-O). These results are consistent with the idea that whereas the ASP cytonemes extend and retract in the HSPG-defective discs at rates that are similar to controls, they had reduced capacity to maintain interactions with cells lacking HSPGs. This idea is consistent both with the steady-state reductions to numbers and lengths of cytonemes in discs that expressed ttvRNAi for at least 24 hours (Fig. 2E-K), and with reduced signaling, since signals transfer between cells at the transient synaptic contacts that ASP cytonemes make with target cells (Huang et al., 2019; Roy et al., 2014).

### Bnl export *ex vivo* and *in vivo*

To investigate the role HS might have in the production and release of Bnl from wing disc cells, we developed a method to monitor Bnl subcellular location. The method is based on imaging GFP fluorescence after GRASP reconstitution of a variant Bnl (Bnl:GFP^11(7x)^) with a split-GFP partner (Feinberg et al., 2008; Feng et al., 2017). The Bnl chimera had an array of seven GFP^11^ sequences (Kamiyama et al., 2016) inserted at a site C-terminal to the FGF domain. Previous work showed that Bnl with an insertion of GFP at this site is functional and is transmitted from disc to the ASP) (Sohr et al., 2019). The GFP partner (GFP^1-10^) was joined to one of several different protein domains in order to concentrate the GFP^1-10^ moiety in a specific subcellular compartment (e.g. the external face of the plasma membrane, endoplasmic reticulum, secretory pathway, or cytoplasm). To verify the subcellular locations of these variants, we analyzed GFP fluorescence in S2 cells that express cytoplasmic GFP as well as either GFP joined to the N-terminus of a CD4-membrane targeting domain (extGFP), to a KDEL-containing ER-targeting domain (erGFP), or to the signal peptide-dependent route in the secretory pathway (secGFP). The fluorescence of extGFP was mostly cell membrane associated, of erGFP in an intracellular distribution consistent with ER-localization, of secGFP in a punctal intracellular distribution, and of cytGFP throughout the cytoplasm (Supplemental Fig. 2A-D). We then analyzed split-GFP fluorescence in S2 cells that express Bnl:GFP^11(7x)^ together with GFP^1-10^ localized to either the plasma membrane (extGFP^1-10^), the endoplasmic reticulum (erGFP^1-10^), or the cytoplasm (cytGFP^1-10^), or traffics by a secretory route (secGFP^1-10^) together with. GFP fluorescence was not observed in the presence of cytGFP^1-10^, but was observed in the presence of extGFP^1-10^, erGFP^1-10^, and secGFP^1-10^ (Fig. 6A-J). These results indicate that Bnl:GFP^11(7x)^ was in an extracellular orientation and GPI-linked in the plasma membrane (Du et al., 2022a), consistent with signal peptide-dependent trafficking.

**Figure 6:**
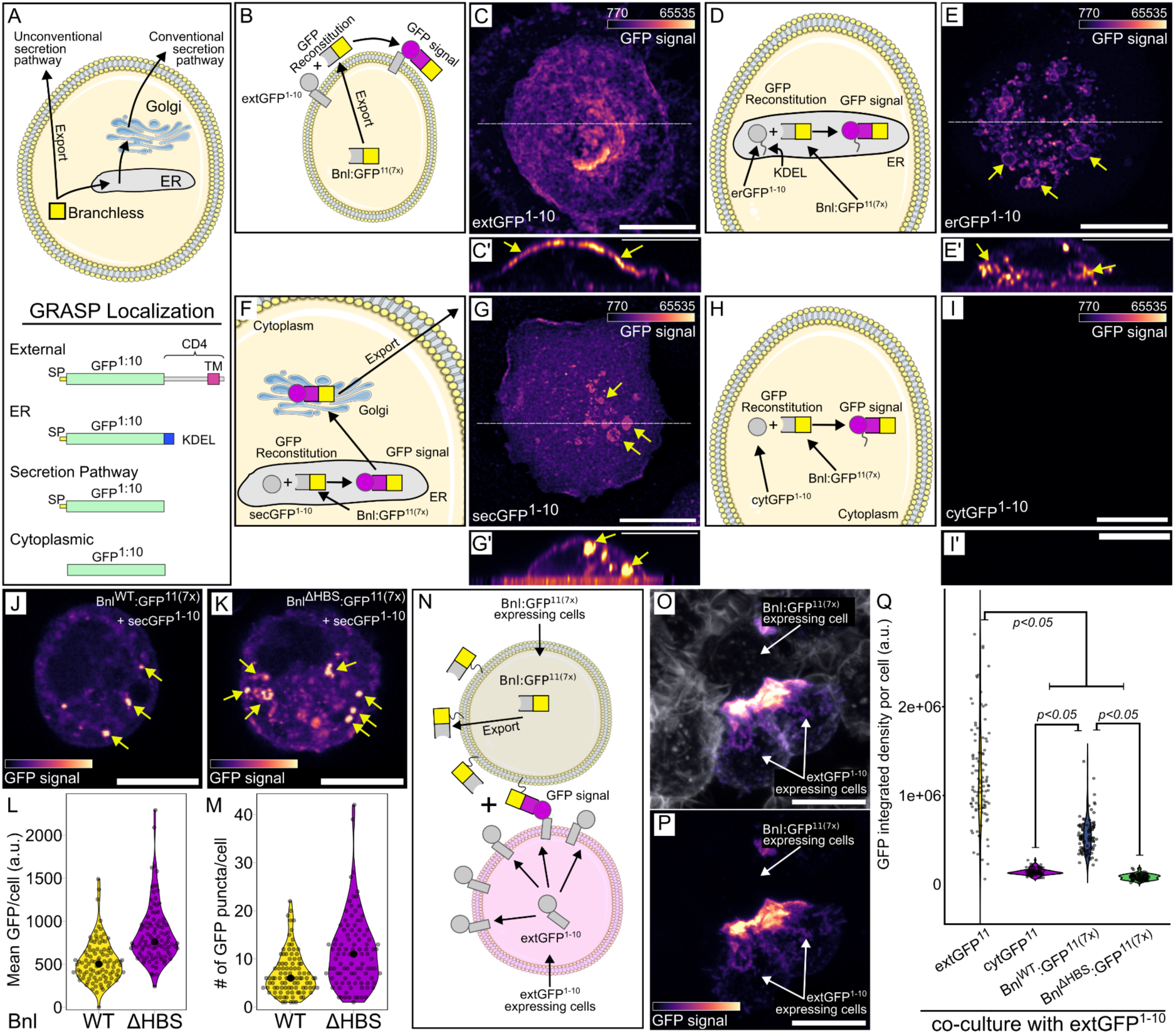
Bnl intracellular trafficking and export. (A) Cartoon depicting Bnl export and GRASP combinations of Bnl:GFP with or without signal peptide (SP), transmembrane domain (CD4) or ER localization tags (KDEL). (B,C,C’) Cartoon depicting SP:GFP^1-10^:CD4 and SP:Bnl:GFP^11(7x)^ GRASP in S2 cells (B) and reconstitution reporting Bnl export to the extracellular side of plasma membrane (C, GFP fluorescence, magenta scale, (C’) yellow arrowheads). (D,E,E’) Cartoon depicting SP:GFP^1-10^:KDEL and Bnl:GFP^11(7x)^ GRASP in S2 cells (D) and reconstituted GFP reporting internal localization (E,E’)). (F,G,G’) Cartoon depicting Sec:GFP^1-10^ and Bnl:GFP^11(7x)^ GRASP in S2 cells (F) and reconstituted GFP fluorescence reporting Bnl export via conventional biosynthetic pathway (G) on the cell membrane and in organelle-like internal structures (G’). (H,I,I’) Cartoon depicting cytoplasmic GFP^1-10^ and Bnl:GFP^11(7x)^ GRASP in S2 cells and reconstituted GFP fluorescence reporting absence of Bnl export through unconventional secretion pathway (I,I’). (pixel intensities in C’,E’,G’,I’ increased to highlight marked structures.) (J-M) Sec:GFP^1-10^ and Bnl:GFP^11(7x)^ GRASP in S2 cells and GFP fluorescence reporting little retention (and export) of WT Bnl (J), but high levels of cytoplasmic Bnl^ΔHBS^. (L,M) Quantification of internal GFP fluorescence of WT Bnl and Bnl^ΔHBS^ (*t*-test, one-tail, *P*<0.001). (N-Q) Cartoon depicting extGFP^1-10^ and Bnl:GFP^11(7x)^ GRASP in separate, co-cultured, S2 cells reporting Bnl-dependent contact, (O,P) S2 cells stained with Alexa Flour^TM^ 647 Phalloidin (white), and reconstituted GFP fluorescence. (Q) Quantification of GFP fluorescence of co-cultured S2 cells expressing either extGFP^1-10^ or extGFP^11(7x)^ (positive control), cytoplasmatic:GFP^1-10^ or extGFP^11(7x)^ (negative control), extGFP^1-10^ or Bnl^WT^:extGFP^1-10^ or Bnl^ΔHBS^ (ANOVA: Df=3, F=310.14. Tukey’s HDS indicated in the plot). (C,E,G,I,O,P) z-stack sum projection, scale bar: 10 µm. (C’,E’,G’,I’) z-orthogonal view, scale bar: 10 µm. (J,K) confocal section view, scale bars: 10 µm.

We next characterized the GRASP fluorescence of Bnl:GFP^11(7x)^ + extGFP^1-10^ in wing discs to determine the degree to which this Bnl construct mimics the normal Bnl protein. The experiment was designed to enhance sensitivity, with the GFP^11^ repeated seven times and with potential aggregation of the GRASP proteins, but we do not have an independent measure of its sensitivity relative to other methods of detection. Understanding that the results are contingent on sensitivity, we expressed Bnl:GFP^11(7x)^ together with membrane-localized mCherry:CAAX in Bnl producing cells (*bnl*-LexA>*lexO*-Bnl:GFP^11(7x)^ *lexO*-mCherry:CAAX), together with CD4-GFP^1-10^ (extGFP) in the dorsal compartment (*ap*-GAL4 UAS-extGFP^11^). In this genotype, all dorsal wing disc cells expressed extGFP^1-10^ and only Bnl-producing cells expressed Bnl:GFP^11(7x)^. As shown in Figure 7(A-E), GFP fluorescence was confined to the domain of Bnl expression. Fewer than 10% of the Bnl-expressing cells had GFP fluorescence; their distribution within the Bnl-expressing domain varied among the discs we examined (Supplemental Fig. 3). The fluorescence was exclusively basal and membrane associated, and was present in a punctal form in some basal cytonemes that extended from the Bnl-expressing cells (Fig. 7D,D’). We did not observe intracellular GFP fluorescence despite the expression of both GRASP partners in every Bnl-expressing cell. This is consistent with the idea that extGFP^1-10^ and Bnl:GFP^11(7x)^ did not associate intracellularly. These results are consistent with the idea that many cells in the Bnl-expression domain make but do not export Bnl at any point in time, and that for a particular cell, secretion is intermittent - that Bnl export is not constitutive and that not all Bnl-expressing cells export Bnl simultaneously.

**Figure 7:**
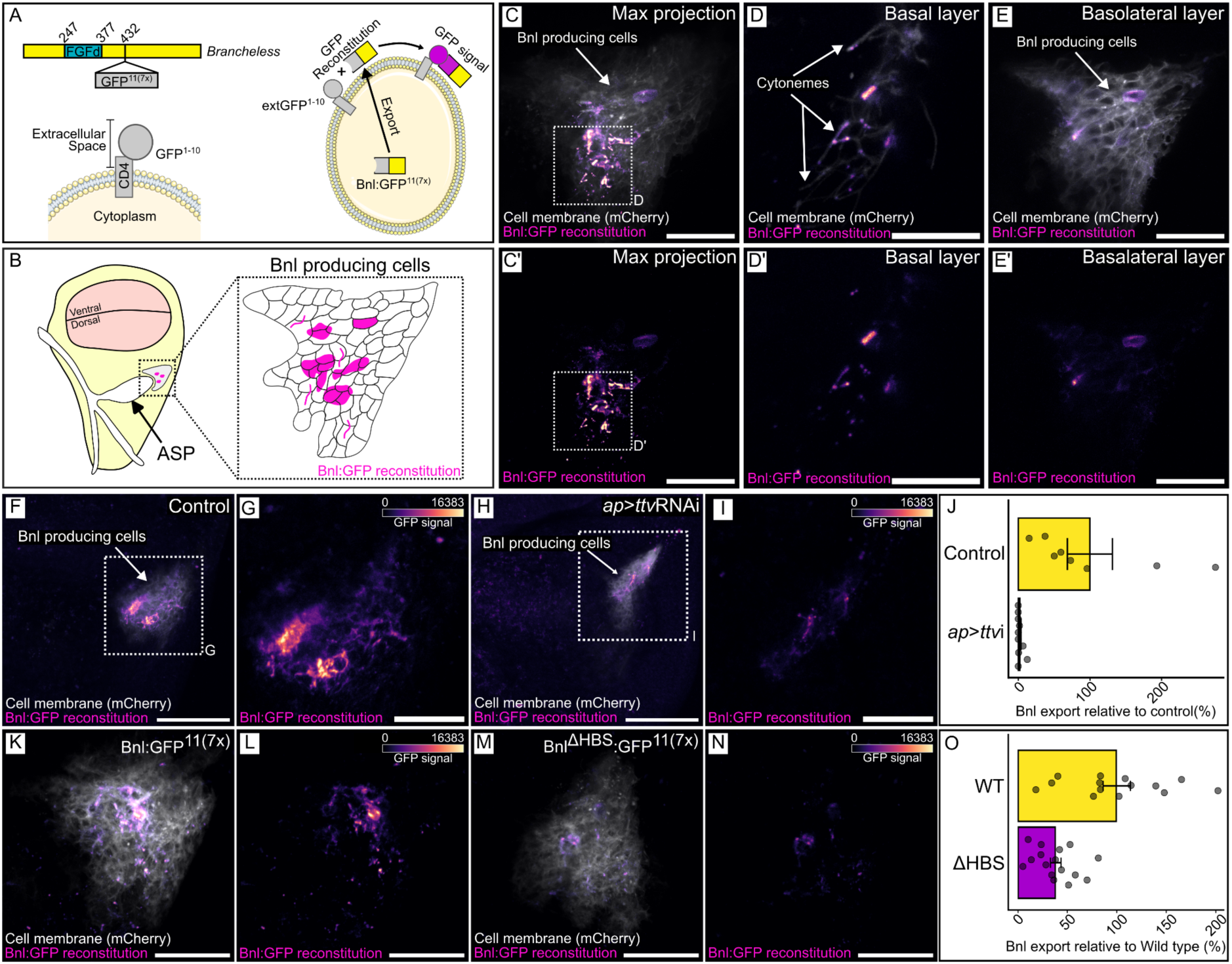
Bnl basolateral localization and presence in cytonemes requires HS. (A) Cartoon depicting Bnl:GFP^11(7x)^ and co-expression with extGFP^1-10^ for GFP reconstitution. (B) Diagram of a wing disc (yellow), illustrating the location of the ASP (white) and the Bnl-producing cells (gray). Inset shows magnified view of Bnl-producing cells, depicting a typical pattern of Bnl:GFP reconstitution (magenta scale). (C-E’) GFP fluorescence (magenta scale) from reconstitution of CD4:GFP^1-10^ expressed in the wing disc dorsal compartment (*ap*-Gal4) and Bnl:GFP^11(7x)^ and Cherry:CAAX (white) expressed in Bnl-producing cells (*bnl*-LexA), showing Max projection images (C,C’), basal layer (D,D’) and basolateral layer (E,E’). (C’,D’,E’) magenta scale only. (F-J) GRASP reconstitution as in (C) in wild type wing disc cells (F,G) or in cells with reduced HS (*ap>*ttvRNAi) (H,I). (G,I) High magnification image of boxed area of (F,H; magenta scale only). (J) Quantification of fluorescence detected in (G,I): (*t*-test, one-tail, *P*<0.05). (K-O) GRASP reconstitution as in (C) with either Bnl:GFP^11(7x)^ or Bnl^ΔHBS^. (L,N) magenta scale only. (O) Quantification of GFP fluorescence detected with either WT (L) or HBS mutant Bnl (N) GRASP. (*t*-test, one-tail, *P*<0.05). Scale bars: (C,C’,E,E’,G,I,K-N), 20 µm, (D,D’) 10 µm, (F,H), 50 µm.

To validate this split-GFP method, we first did a control experiment with the same general setup except we expressed extGFP^1-10^ together with extGFP^11^ in place of Bnl:GFP^11(7x)^. Membrane-associated GFP fluorescence was observed throughout the Bnl domain, but not elsewhere in the dorsal domain (Fig. 8A,B). The GRASP fluorescence of these constructs was not dependent on HS production (Fig. 8C,D). These results are consistent with the idea that the method we used marks all cells that have a split-GFP partner for extGFP^1-10^ on their surface, and that only some of the cells had sufficient Bnl:GFP^11(7x)^ to generate GRASP fluorescence with extGFP^1-10^.

**Figure 8:**
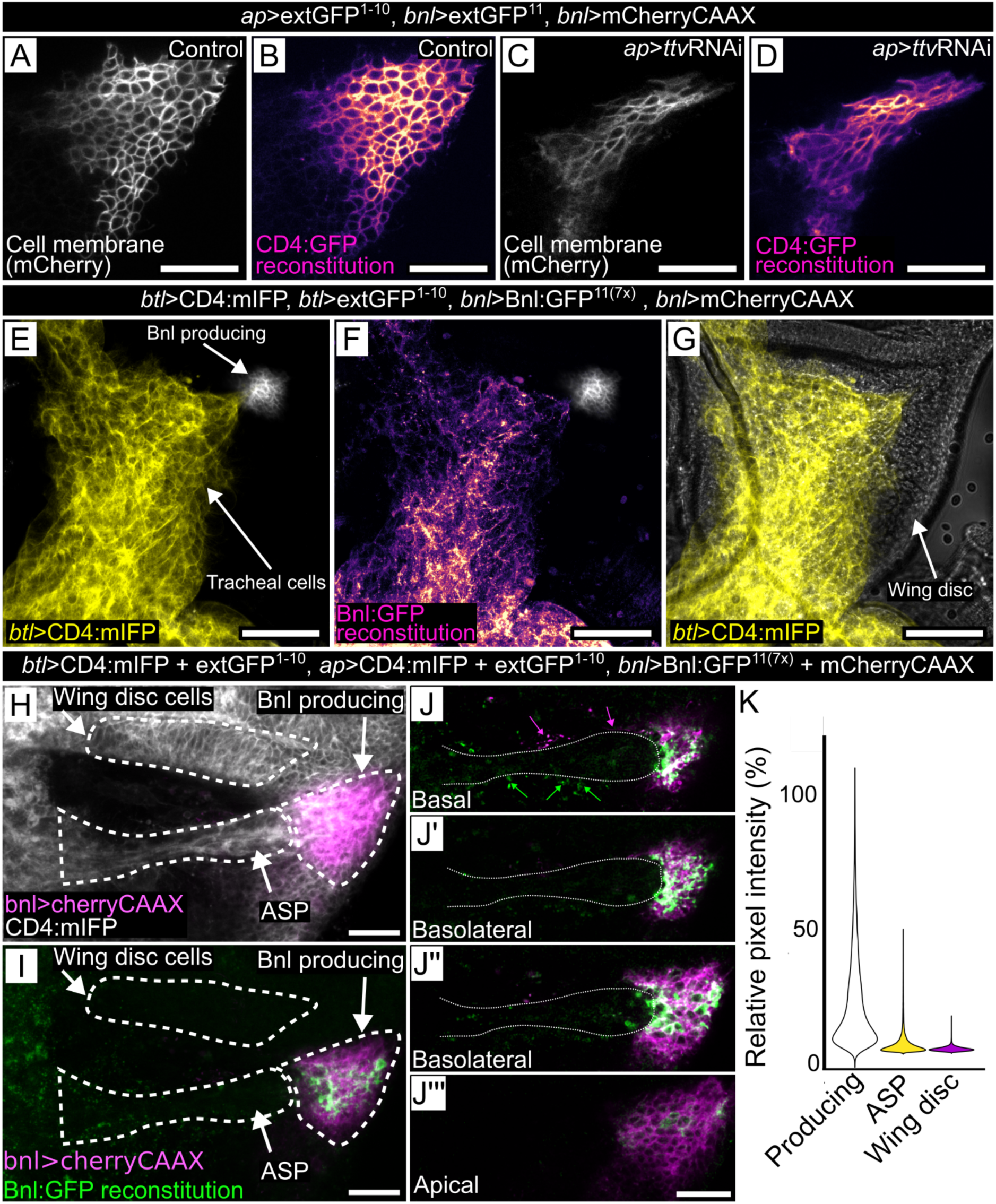
Bnl:GFP^11(7x)^ is an active signaling protein. (A-D) Control (A,B) and ttvRNAi knockdown conditions for GFP reconstitution of ext:GFP^11^ produced by Bnl-expressing wing disc cells (*bnl*-LexA lexO-CD4:GFP^11^) and ext:GFP^1-10^ produced by dorsal wing disc cells (*ap*-Gal4 UAS-CD4GFP^1-10^) in membranes of Bnl-expressing cells. (A,C) membranes marked with mCherry:CAAX fluorescence (white); (B,D) membranes marked with GFP fluorescence (magenta scale). (C,D) ttvRNAi expressed by UAS-ttvRNAi transgene. (E-G) Aberrant growth of the ASP and tracheal branch, marked with cell membrane protein CD4:mIFP (E,G - yellow) and (F) Bnl:GFP fluorescence of reconstituted GFP. Bnl:GFP^11(7x)^ over-expression by Bnl producing cells (*bnl*-LexA) caused overgrowth of cells expressing extGFP^1-10^ (*btl*-Gal4, UAS-CD4:mIFP (yellow)). (H) Bnl:GFP^11(7x)^ over-expression by Bnl producing cells (*bnl*-LexA) together with CD4:GFP^1-10^ and CD4:mIFP expressed in the wing disc dorsal compartment (*ap*-Gal4) and the ASP (*btl*-Gal4) did not induce ASP overgrowth. (I) GFP reconstitution (green) was mostly on Bnl-producing cells (magenta). (J) Confocal sections showing Cherry-CAAX (magenta) in Bnl-producing cells and extending along the ASP (magenta arrows), and GFP reconstitution (green) at the surface of Bnl-producing cells and extending along ASP (green arrows). (K) Pixel intensity plots of fluorescence in the boxed areas of Bnl-producing and other disc cells, and ASP, relative to maximum value. Scale bar: (A-G) 50 µm, (H,I) 20 µm.

Another possible explanation is that Bnl:GFP^11(7x)^ export is constitutive, but that its release or removal from the cell surface is more efficient than binding to the split-GFP partner. We examined this possibility by first analyzing the biological activity of Bnl:GFP^11(7x)^. We expressed Bnl:GFP^11(7x)^ in the Bnl domain of the wing disc and expressed extGFP^1-10^ in the ASP (*btl*-Gal4 UAS-extGFP^1-10^ UAS-mIFP *bnl*-LexA>*lexO*-Bnl:GFP^11(7x)^ *lexO*-mCherry:CAAX). GFP fluorescence was both membrane-associated and punctal (Fig. 8E-G), and expression of the Bnl:GFP^11(7x)^ led to tracheal overgrowth and no apparent ASP, a phenotype that is indistinguishable from the effects of ectopic over-expression of Bnl:GFP in the ASP (Sohr et al., 2019). This result is consistent with the idea that the Bnl:GFP^11(7x)^ has activity that is comparable to Bnl:GFP and that the complex with extGFP^1-10^ forms efficiently and can activate FGF signal transduction to levels comparable to Bnl:GFP, despite localization of the GRASP-tethered Bnl to the membrane of a FGFR-containing cell.

We also set up a competitive assay by expressing extGFP^1-10^ in both the ASP and wing disc and expressing Bnl:GFP^11(7x)^ in the Bnl domain of wing disc (*btl*-GAL4 *ap*-GAL4 UAS-extGFP^1-10^ *bnl*-LexA *lexO*-Bnl:GFP^11(7x)^). In contrast to discs in which extGFP^1-10^ was expressed only in the ASP, no ASP overgrowth was observed. Moreover, despite the presence of extGFP^1-10^ throughout the wing disc dorsal compartment, fluorescence of reconstituted GFP was only observed on the surface of wing disc cells in the Bnl domain (Fig. 8H-J). These results are consistent with the idea that Bnl:GFP^11(7x)^ is active and signals to tracheal cells, but that reconstitution with membrane-tethered GFP captured it on cell surface and reduced its potency as a paracrine signal.

### Role of HS in Bnl export

To determine if the split-GFP system might also be used to better characterize the role of HS in Bnl export, we expressed Bnl:GFP^11(7x)^ in Bnl producing cells of the wing disc and extGFP^1-10^ and ttvRNAi in the wing disc dorsal compartment (*bnl*-LexA>*lexO*-Bnl:GFP^11(7x)^ *ap*-GAL4 UAS-extGFP^1-10^ UAS-ttvRNAi). GFP fluorescence was reduced by ∼98% in HS-depleted discs (Fig. 8F-J), consistent with inhibition of Bnl export and with the previous experiments showing reduced ASP development in discs that were depleted of HSPGs by expression of ttvRNAi (Fig. 1 and 2). We next set up contexts which inhibit HS binding to Bnl in order to better understand the role HS has as a component of an HSPG.

As noted in the Introduction, HS binding to a FGF HBS is driven by electrostatic interactions between positively charged arginine and lysine residues in the HBS and negatively charged sulfate groups of HS (Xu and Esko, 2014). In order to reduce the affinity of the Bnl for HS, we identified a candidate HBS in Bnl and changed arginine and lysine residues in the HBS to glutamate. The residues were selected by first generating a putative structure with Alphafold (Jumper et al., 2021) to identify the 12 stranded β-trefoil domain characteristic of the conserved FGF domain (residues T247-I377), then performing a heparan docking analysis with ClusPro (Kozakov et al., 2017), and lastly identifying the residues with the highest contact frequency and lowest negative interaction energy (K256, R343, R344, R357, R358, R365, K369) using the Molecular Operating Environment (MOE) program (Fig. 9A-C). These seven residues are predicted to be on the surface of the FGF domain and to have the highest potential to interact with HS. We also mutated K258 for this study because of its location in the putative HBS and because of its orientation predicted by the docking analysis, despite the finding that its contact frequency and interaction energy with HS were not significant. The Bnl variant with K256E, R258E, R433E, R434E, R357E, R358E mutations (Bnl^ΔHBS^) was characterized for this study. The electrostatic potential of its HBS is expected to be reversed, and molecular dynamics analysis indicates that the binding energy of the mutant FGF domain with HS is significantly reduced (Fig. 9D-I). We used the split-GFP system for subcellular localization analysis to characterize the behavior of Bnl^ΔHBS^ mutants in both cultured cells and wing discs.

**Figure 9:**
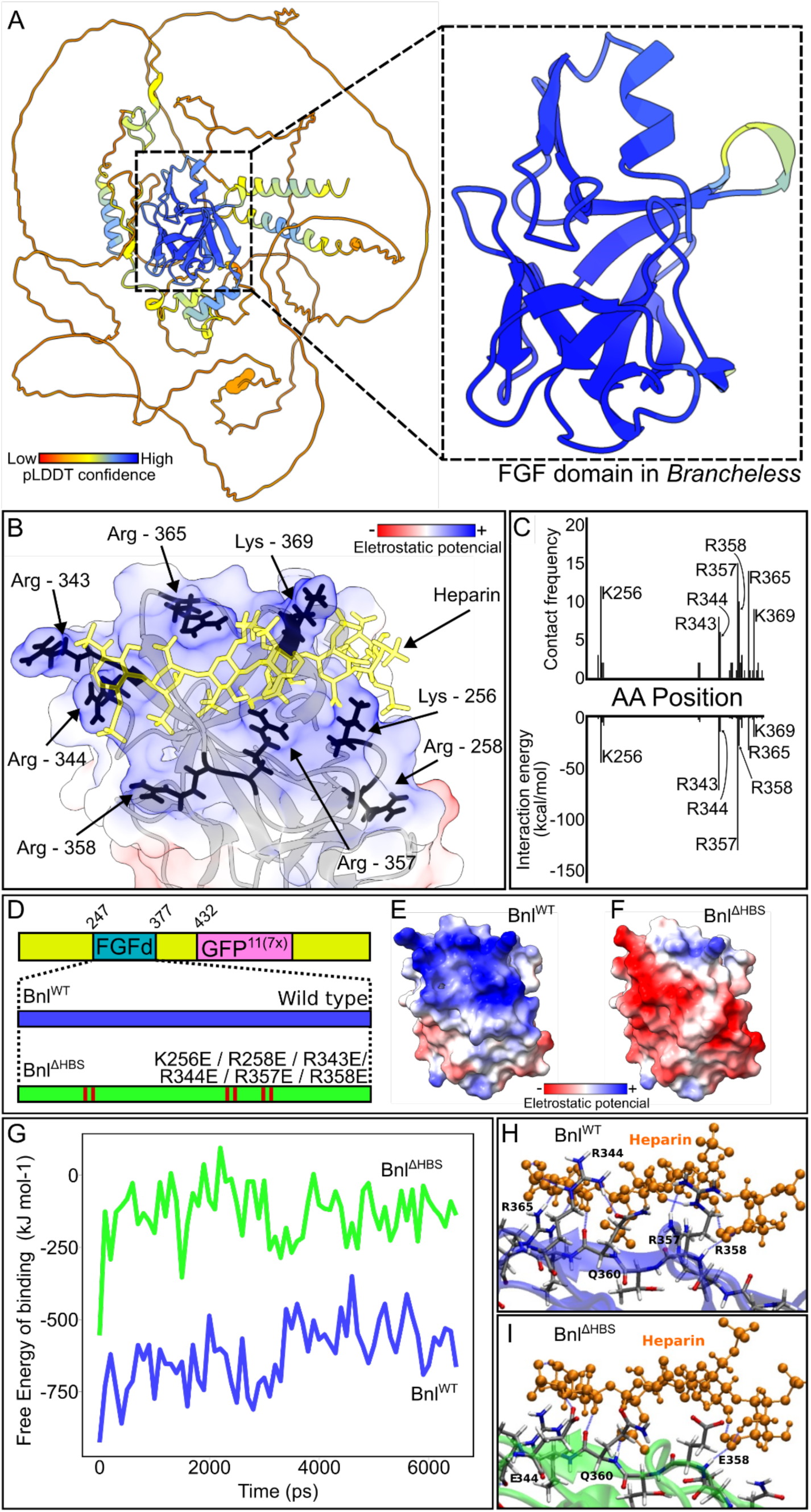
Structure-guided design of mutant Bnl HBS. (A) Predicted Alphafold structure of Bnl has a β-trefoil FGF domain composed of 12 β-strands (dashed box). (B) Eight lysine and arginine residues (black) predicted by ClusPro to form an HBS surface with positive electrostatic potential. (C) K265, R258, R343, R344, R357, R358, R356, and K369 predicted to contact HS at higher frequency and lower interaction free energy during Bnl and heparin docking. (D) Structure-guided mutations in the FGF domain, generating the HBS variant (ΔHBS) replacing the target lysines (K) or arginines (R) with glutamicate (E), changing the electrostatic potential from (E) positive (blue) to (F) negative (red). (G) Free energy of binding estimated on molecular dynamic simulation of heparin binding with (H) Bnl^WT^ and (I) or Bnl^ΔHBS^.

In S2 cells that express sec:GFP^1-10^ and either Bnl^WT^:GFP^11(7x)^ or Bnl^ΔHBS^:GFP^11(7x)^, the fluorescence of the Bnl^WT^ construct was extracellular, in contrast to the fluorescence of the Bnl^ΔHBS^ construct, which was intracellular in these confocal optical sections, and more intense (Fig. 6J-M). We suggest that a likely possibility is that the complex of WT Bnl and secGFP was secreted but that the complex with the mutant Bnl^ΔHBS^ was not. We also generated S2 cells that express either Bnl^WT^:GFP^11(7x)^ or Bnl^ΔHBS^:GFP^11(7x)^ and co-cultured them with S2 cells expressing extGFP^1-10^. For the combination with Bnl^WT^:GFP^11(7x)^, GFP fluorescence was observed at regions of contact between cells, similar to the positive control (extGFP^11^ and extGFP^1-10^), but the intensity of fluorescence with Bnl^ΔHBS^:GFP^11(7x)^ was lower, and not detectably greater than a negative control (cytGFP^1-10^ and extGFP^11^ cells) (Fig. 6N-Q). These results are consistent with the idea that Bnl protein that does not bind HS is not exported efficiently and instead accumulates inside the cell.

To confirm that these results reflect *in vivo* behavior, we expressed Bnl^ΔHBS^ in wing discs and monitored Bnl^ΔHBS^ distribution with the split-GFP system. We compared GFP fluorescence in discs that express extGFP^1-10^ in the dorsal compartment (*ap*-GAL4 UAS-extGFP^1-10^) and express either Bnl^WT^:GFP^11(7x)^ or Bnl^ΔHBS^:GFP^11(7x)^ in Bnl-producing cells (*bnl*-LexA lexO-Bnl^WT^:GFP^11(7x)^ or *bnl*-LexA lexO-Bnl^ΔHBS^:GFP^11(7x)^). As described above, whereas a small number of Bnl-expressing cells had Bnl^WT^ on their surface (Bnl^WT^:GFP^11(7x)^ + extGFP^1-10^), and GFP fluorescence was reduced by more than two-thirds with the mutant Bnl^ΔHBS^ (Bnl^ΔHBS^:GFP^11(7x)^ + extGFP^1-10^) (Fig. 7K-O).

## Discussion

Although HS is known to be essential for FGF signaling due to the participation of an HSPG as co-receptor, HSPGs are likely to have additional roles in FGF signaling. FGF signaling involves many discrete steps, including synthesis, processing, intracellular transit, export to the cell surface, and release to the FGF receptor. In the Drosophila system we investigated, FGF signaling is dependent on Dlp which is produced by the wing disc (Du et al., 2018; Huang and Kornberg, 2016; Sato and Kornberg, 2002), but it is not known specifically what Dlp does or what other roles HSPGs might have in the disc cells that make Bnl, in the disc cells situated between the cells that make Bnl and the ASP, or in the ASP cells that have the Btl FGFR and take up Bnl. The studies we report here identify a role for HS in Bnl-expressing disc cells and, as previously proposed by Du et al (2018), a role of surface Bnl for the ASP cytonemes that carry Bnl to the ASP. Our unexpected observation that only a minority of Bnl-producing disc cells had detectable Bnl on their surface has intriguing general implications for the mechanisms that regulate export and dissemination of signaling proteins.

Studies of Drosophila mutants deficient for either HS or for the protein core of an HSPG have reported defective signaling by Hedgehog (Hh), Decapentaplegic (Dpp), Wingless (Wg), and FGF (Bilioni et al., 2013; Bischoff et al., 2013; Han et al., 2004; Lin et al., 1999; Takei et al., 2004). Given the prevailing model - that these proteins move through tissues by passive “restricted” diffusion (Li et al., 2018) - the inability of Hh, Dpp and Wg to cross efficiently over HS-deficient cells was interpreted to be a consequence of their inability to bind HSPGs linked to cell surfaces or to the associated ECM. From our perspective, it is now established that these signaling proteins move across tissues linked to cytonemes and transfer directly from a producing to a target cell at cytoneme synapses, and it is therefore imperative to understand how HS deficiencies affect cytoneme-mediated dispersion. The following discussion offers our perspective based on the fact that Bnl signaling is dependent on release of GPI-linked Bnl from disc cells at cytoneme synaptic contacts (Du et al., 2022a, 2018) and on cytoneme transport from the cytoneme synapse to the recipient ASP cell.

Bischoff et al (2013) reported that in fixed preparations, cytonemes required for Hh signaling were not observed extending over *ttv botv* mutant cells deficient for HS, and suggested that the ECM overlying mutant cells might be compromised and unable to stabilize cytonemes (Bischoff et al., 2013). The conditions we tested (RNAi-mediated inhibition) also reduced HS (Fig. 1) and impaired signaling (Fig. 2), but the evidence we obtained does not support the idea that the ECM was compromised (Fig. 5). Instead, imaging active cytonemes in unfixed preparations revealed that although cytonemes were observed to extend over HS-deficient cells, their length, lifetime and stalling time were reduced (Fig. 4). We suggest that these features may indicate that cytonemes extend normally over HS-deficient cells, but their reduced capacity to make functional contacts led to the observed effects on cytoneme behavior and signaling. Similar behaviors have been reported previously for cell extensions in other mutant contexts.

In the Drosophila visual system, axons extending from R8 ommatidial neurons appear to fail to reach their targets in mutants that lack Netrin or the Netrin receptor Frazzled (Timofeev et al., 2012). Although it was initially assumed that this abnormality was due either to the absence of a chemoattractant gradient of extracellular Netrin, or to lack of response to a Netrin gradient, live imaging revealed otherwise (Akin and Zipursky, 2016). The growth cone filopodium of the R8 axon was observed to extend to its target in both normal and mutant fly brains, and in normal brains the contact was stable and led to a functional synapse. In contrast, contact with the target was transient in the mutants, and the subsequent retraction of the mutant growth cone gave the misleading appearance of its inability to reach the target. Moreover, the effects of HS deficiency in mouse commissural neurons phenocopies Netrin mutants (Matsumoto et al., 2007), evidence for a cell autonomous HS function that is similar to the intracellular role we observed for Drosophila Bnl.

HSPGs are also needed by the dendrites of space-filling C4da peripheral neurons that innervate the Drosophila larval skin (Poe et al., 2017). Dendrite growth in areas of normal or HSPG-deficient epidermis was indistinguishable, but in contrast to normal animals, dendrite density and arborization was not maintained in HSPG-deficient areas. The authors determined that destabilization of the dendritic arbors was not due to failure to bind a known HSPG receptor on the neuron, or to failure of HSPG-mediated transport of a putative diffusible ligand. Instead, their findings implicated a receptor phosphatase and a mechanism that involves direct dendrite-epidermal cell contact. They suggest that their results are consistent with the idea that HS-deficient epidermal cells fail to present an essential ligand to the dendrite. We speculate that the ligand may be Maverick (Mav), a member of TGF-β superfamily produced by the epidermis (Hoyer et al., 2018) and that activates the Ret receptor of C4da dendritic projections (Soba et al., 2015). Most TGF-β superfamily proteins bind HSPGs, and it is possible that export of Mav to the cell surface may be HSPG-dependent.

A landmark study of Drosophila Hh from the Guerrero lab reported an intracellular role for an HSPG in paracrine signaling (Callejo et al., 2011). Newly synthesized Hh was found to attach first to the apical plasma membrane before internalization in vesicles that move intracellularly to basal cytonemes that transport Hh to target cells. Endocytic uptake of Hh from the apical membrane and apical to basal trafficking of Hh-containing vesicles was impaired in Dlp mutant cells. Subsequent studies reported that HSPGs were also required for the Hh co-receptor Ihog to be detected at the cell surface (Bilioni et al., 2013; Simon et al., 2021), and that HS mediates a high affinity interaction between Hh and Ihog in a tripartite Ihog:Hh:HSPG complex that is essential for the cytonemes that generate the Hh concentration gradient (McLellan et al., 2006; Simon et al., 2021). In mouse, HSPGs have also been found to be essential for the gradient distributions of FGF7 and FGF10 (Makarenkova et al., 2009). Together with our observation that the levels of cell surface Bnl were reduced in conditions that eliminate HS or abolish HS:Bnl binding (Figs. 6,7), these findings are consistent with the idea that direct interactions with HS are essential for the processes that move signaling proteins to the cell surface.

In the wing disc system that includes the disc epithelium, hemocytes, myoblasts, and tracheal cells, only tracheal cells produce the Btl FGFR, and Bnl production is limited in both space and time (Sato and Kornberg, 2002). We understand that transcriptional regulation specifies the place and time of Bnl and Btl production (Ohshiro et al., 2002), but there is much evidence for additional regulatory processes that control how Bnl disperses from the disc cells to engage Btl in the trachea. One that has been identified by the Roy lab is a system of positive feedback that responds to Bnl signal transduction to determine the number of cytonemes that extend between the ASP and the disc (Du et al., 2018). A key attribute of this system is that it defines the tissue-specific contours of Bnl distributions in real time by suppressing the number of cytonemes extending from cells with low levels of signal transduction and increasing the number of cytonemes extending from cells with higher levels of signal transduction. Bnl secretion is dependent on Furin-dependent pro-domain cleavage and GPI-linkage, which promotes trafficking to basal membranes, where ASP cytonemes pick up Bnl (Sohr et al., 2019).

Subsequent studies showed that contact-mediated Bnl release involves responses in both the Bnl-producing disc cell and the receptor-bearing ASP recipient cell such that source and recipient cytonemes reciprocally guide each other to form signaling contacts.(Du et al., 2022b). This complex but still undefined process provides ways to regulate both selectivity and levels, and its existence is consistent with our finding that ASP cytonemes that mediate Bnl signaling to the ASP do not behave normally in areas with reduced levels of cell surface Bnl.

There is also evidence that release of Hh, Wg, and Dpp is regulated at cytoneme synapses (Hatori et al., 2021). Whereas the amount of Hh mRNA and Hh protein produced in the wing disc is proportional to gene copy number, the amount of Hh protein released to target cells is not: the amount of Hh observed in target cells is the same in animals with 1, 2, 3, or 4 Hh genes. Wg and Dpp signaling are similarly insensitive to levels of production. These properties are reminiscent of neurotransmitter release that is regulated at neuronal presynaptic terminals. Cytoneme synapses may be similarly programmed such that exchanges of signaling proteins are initiated only upon stimulation.

Our finding that the presence of detectable Bnl on the cell surface depends on a process that requires HS is consistent with the idea that Bnl export is not constitutive. Precedents for an intracellular HSPG function is provided by the Guerrero studies of Hh signaling (Callejo et al., 2011) and by a study showing that HSPGs are essential for the export of vertebrate Sonic Hh from the Golgi (Tang et al., 2022). Bnl and Hh are signaling proteins that contribute to growth control, and it is interesting to consider what factors might regulate the intracellular trafficking pathway that controls their release. Our findings show that the HSPGs might be essential components of this pathway.

## Acknowledgements

We thank members of the Kornberg lab for their help: Sol Fereres Rapoport, Ryo Hatori, Huang Hai, Brent Wood, Wanpeng Wang, Zehra Ail-Murthy, Will Fleming, Songmei Liu. We thank Daniel R. Sandoval for assisting with the Molecular Operating Environment (MOE) analysis and the Bloomington Drosophila Stock Center (NIH P40OD018537) for fly stocks. Funding: NIH GM122548

## Materials and Methods

### HS Biosynthesis gene expression

Two pools containing 10-15 wing imaginal discs of wild type L3 larvae were dissected in PBS and transferred to 300 µL RTL Buffer (Qiagen RNeasy Micro Kit). RNA extraction was performed according to manufacturer instructions. cDNA was synthesized using SuperScript™ III Reverse Transcriptase kit (Invitrogen) with oligo dT according to manufacturer instructions. qRT-PCR was performed on a C1000 Touch PCR thermal cycler (BioRad) using SYBR Green (Bioline). Efficiency of each primer pair was determined by standard dilution analysis (1:2) with wing disc cDNA. Specificity of each primer was determined by standard melting curve analysis after PCR cycles. Primers were designed using Primer-BLAST (NCBI) following instructions for SYBR Green detection.

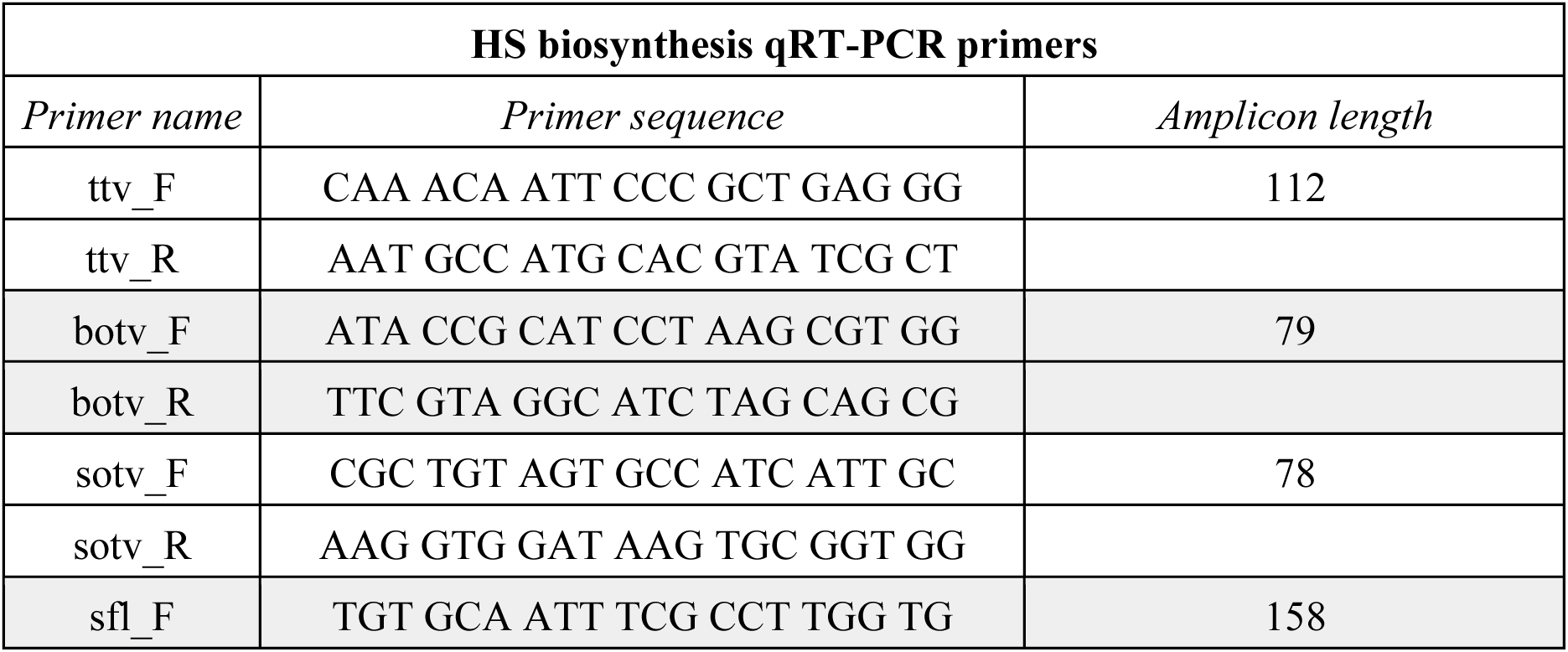
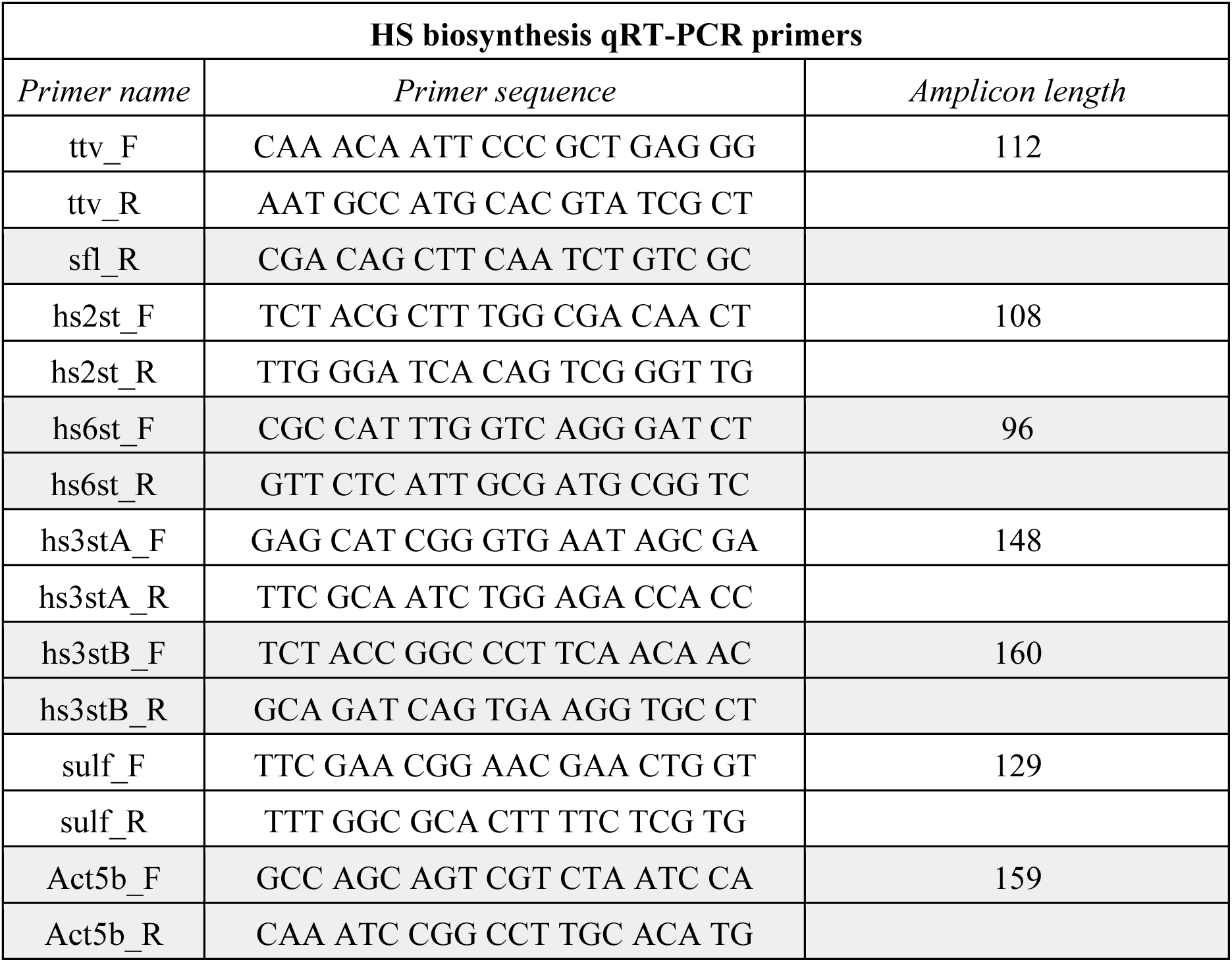

### Drosophila stocks and husbandry

Flies were cultured on standard cornmeal and agar medium at 25°C except for temporal control of *ttv* RNAi expression: incubation was at 18°C prior to temperature shift to 29°C for inactivation of GAL80^ts^ for 24, 48 and 72 hours and collection of wandering stage L3 larvae. For ASP morphogenesis analysis, *ap*-Gal4/CyO,Wee-P;Btl-LHG,LexO-CD4:GFP flies were crossed to UAS-RNAi flies for *ttv, botv, sotv, slf, hs2st, hs6st, hs3sta, hs3stb* and *sulf*, and to UAS-SULF1 for Sulfatase1 overexpression. For time lapse analysis, *btl*-LHG,LexO-CD2:GFP/CyO,Wee-P;*hh*-GAL4,UAS-CD4:mIFP/TM6b was crossed either with w- (control) or UAS-ttvRNAi/CyO,Wee-P. For Bnl HBS studies, *ap*-GAL4/CyO,Wee-P; Bnl-LexA, LexO-mCherry-CAAX flies were crossed with UAS-CD4:GFP^1-10^, LexO-Bnl(ΔHBS variants):GFP^11(7x)^. *ap*-GAL4/CyO,Wee-P; Bnl-LexA, LexO-mCherry-CAAX were also crossed with UAS-ttvRNAi/CyO,Wee-P ; UAS-CD4:GFP^1-10^ LexO-Bnl:GFP^11(7x)^/TM6b.

### Immunostaining

The anterior half of L3 wandering stage larvae was dissected and the cuticle was flipped inside-out for removal of fat body, gut, and ventral nerve cord, leaving attached imaginal discs and trachea. Fixation was in 4% paraformaldehyde for 20 min, and after rinsing in PBS, specimens were permeabilized with PBST (PBS + 0.3% TritonX-100) for 10 min, blocked for 1h with PBST+3%BSA (blocking buffer), and incubated with primary antibodies overnight at 4°C. Specimens were extensively washed with PBS-T and incubated with secondary antibody in PBS for one hour at room temperature in darkness, washed with PBS and transferred to a 20µL PBS droplet in a slide where the wing discs were isolated. PBS was removed and samples were mounted in Vectashield and imaged on Olympus FV3000 inverted laser scanning confocal microscope using an oil immersion objective, UPLFLN40XO.

| Primary antibodies |
| --- |
| dpERK - Phospho-p44/42 MAPK (Erk1/2) (Thr202/Tyr204) (D13.14.4E) Rabbit Monoclonal Antibody #4370 Cell Signaling Technology |
| Patched - Mouse monoclonal anti-Patched (extracellular region) antibody (DSHB Cat# Drosophila Ptc, RRID:AB_528441) |
| Cut/myoblast - Mouse monoclonal anti-Cut homeobox antibody (DSHB Cat# 2b10, RRID:AB_528186) |
| Secondary antibodies |
| Goat anti-Rabbit IgG (H+L) Cross-Adsorbed Secondary Antibody, Alexa Fluor™ 488 Invitrogen |
| Goat anti-Rabbit IgG (H+L) Cross-Adsorbed Secondary Antibody, Alexa Fluor™ 555 Invitrogen |
| Goat anti-Mouse IgG (H+L) Cross-Adsorbed Secondary Antibody, Alexa Fluor™ 488 |
| Goat anti-Mouse IgG (H+L) Cross-Adsorbed Secondary Antibody, Alexa Fluor™ 555 Invitrogen |

### Cytoneme live imaging

Live imaging of the cytonemes was performed as previously described (Barbosa and Kornberg, 2022). Briefly, the wing disc tethered to its tracheal branch were dissected from L3 wandering larvae in Schneider’s media (Gibco; Cat#:21720024) supplemented with 2%–5% FBS. The sample was gently placed on the glass surface of a 35 mm Glass Bottom μ-Dish (Ibidi, Cat#: 81158). The sample was held next to the glass by the placement of a hydrophilic Membrane Filter (Millipore HAWP01300) either with the assistance of double sided tape or a 3D printed ring device (Barbosa and Kornberg, 2022), and 2 mL of medium was added to the dish. Z-stack images were obtained on a Olympus FV3000 inverted laser scanning confocal microscope for one hour at one frame per minute (i.e.: 60 stacks with 0.5 μm spacing imaged at 1 stack/min). After acquisition, images were denoised using Noise2Void (2D) using GoogleColab scripts by ZeroCostDL4Mic (von Chamier et al., 2021).

### Morphometrics measurements on FIJI/ImageJ

Calibrated microscope images were used for measurements using FIJI/ImageJ (Schindelin et al., 2012). ASP length was measured as the distance from the transversive connective to the ASP distal tip; width was measured as the largest perpendicular distance in the ASP bulb. The wing disc dorsal compartment width was the measured length of a line drawn along the ASP long axis from the anterior to posterior edge of the disc. Width measurements of wing disc A and P compartments were made by tracing a line along the dorsal/ventral compartment border from the A/P border to the edge of the measured compartment. Cytoneme counts, and static and dynamic length measurements were assisted by the Cytoneme Analysis tool on FIJI (Barbosa and Kornberg, 2021). For LanA:GFP fluorescence measurements, the confocal z-stack image was sum projected and the integrated density pixel in the dorsal compartment area was measured for each sample and the final result of each sample was plotted relative to the control average measurements.

### Branchless heparin binding site prediction

Banchless alphafold predicted structure (Jumper et al., 2021) was assessed with ChimeraX (Uniprot-id: Q9VDT9) (Pettersen et al., 2021). The FGF domain ranging from amino acids T247 to I377, was filtered and saved as a PDB file. The Bnl FGF domain docking with heparin was performed on ClusPro (Kozakov et al., 2017) and the lysine/arginine residues with most heparin predicted contact were listed for investigation of the Bnl FGF domain Heparin Binding Site (HBS). Finally the models from ClusPro were analysed in Molecular Operating Environment (MOE) software to identify heparin-protein contacts and energy contributions.

### Bnl:GFP^11(7x)^ design and ΔHBS mutagenesis

The Bnl was inserted into a pAc5.1/V5-His A plasmid (ThermoFisher Cat#:V411020) with a sequence of GFP_11_ along with a 5 residue spacer repeated 7 times (Kamiyama et al., 2016). GFP^11(7x)^ was inserted between residues 432 and 433, located C-terminal to the FGF domain of Bnl (Sohr et al., 2019), using Gibson Assembly Cloning Kit (NEB Cat#: E5510S), Q5® High-Fidelity DNA Polymerase (NEB #M0491) and pAc5.1/V5-His A linearized by XhoI digestion.

Based on the ClusPro findings, a set of eight primers were designed to create four Bnl^ΔHBS^ variants with mutations replacing the lysine and arginine residues with glutamic acid. The mutations were made on pAc5.1-Bnl:GFP^11(7x)^ with the Q5® Site-Directed Mutagenesis Kit (NEB Cat#: E0554S).

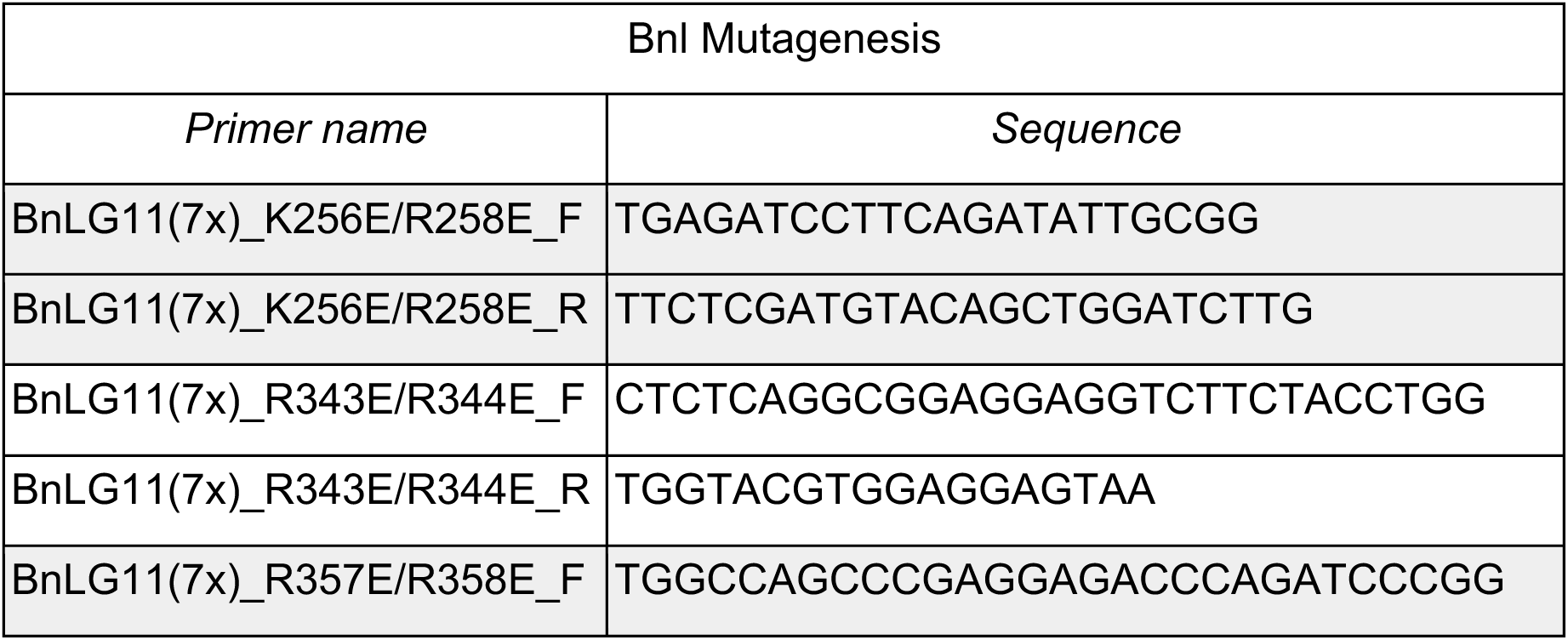
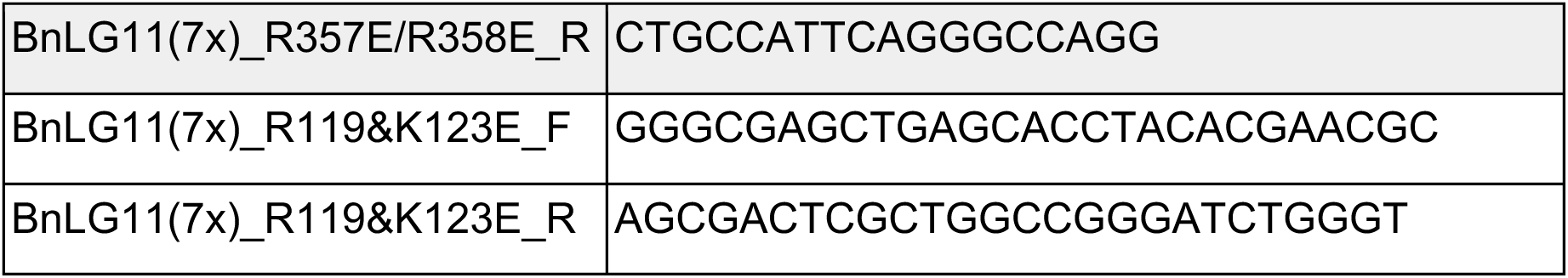

A set of four plasmids were created to track the biosynthetic pathway of Bnl by GFP reconstitution. First, the construct pAc5.1-SignalPeptide-GFP^1-10^ was derived from pUASt-GFP^1-10^-CD4 after cloning GFP^1-10^-CD4 into pAc5.1 followed by deletion of the CD4 sequence to create pAc5.1-SP-GFP^1-10^. pAc5.1-SP-GFP^1-10^-KDEL was generated from pAc5.1-SP-GFP^1-10^ by deleting the signal peptide sequence and attaching KDEL.

To isolate transgenic flies, the plasmid pJFRC19-13XLexAop2-IVS-myr::GFP (Pettersen et al., 2021b) was digested with XhoI and XbaI (removing the myristoylated, codon-optimized GFP) and the wild type and ΔHBS variants were inserted into XhoI and XbaI cloning site, creating the LexOp-Bnl:GFP^11(7x)^ variant plasmid. The pJFRC19-13XLexAop2-Bnl:GFP^11(7x)^ plasmids were modified by adding a UASpromoter-SignalPeptide-GFP^1-10^-CD4-SV40-PA-Terminator sequence into a HindIII restriction site via Gibson Assembly. The final plasmids resulted in an attB-containing pJFRC19-13XLexAop2-Bnl:GFP^11(7x)^/UASt-SignalPeptide-GFP^1-10^-CD4 for each Bnl variant. All plasmids were verified by sequencing and injected together with phiC31 plasmid. All inserted constructs were on the third chromosome of y[1] w[1118]; PBac{y[+]-attP-3B}VK00033 (BDSC Stock#9750).

### Bnl:GFP^11(7x)^ localization assay in S2 Cells

S2 cells were cultured in Schneider’s media supplemented with 5% FBS. Cells were transfected using TransIT®-Insect Transfection Reagent (SKU: MIR 6105). The pAc5.1-Bnl:GFP^11(7x)^ plasmid was co-transfected with either pAc5.1-SP-GFP^1-10^-CD4, pAc5.1-SP-GFP^1-10^, pAc5.1-SP-GFP^1-10^-KDEL or pAc5.1-GFP^1-10^. For cell contact Bnl transfer experiments, S2 cells were transfected with pAc5.1-Bnl:GFP^11(7x)^ or with pAc5.1-SP-GFP^1-10^-CD4. 48 hours after transfection, both cell populations were collected, washed with PBS, and co-cultured in a well for 24 hours.

Positive controls for this experiment were experiments with one population transfected with pAc5.1-SP-GFP^1-10^-CD4 and the other pAc5.1-SP-GFP^11^-CD4. Negative controls were with cells transfected with either pAc5.1-cytoplasmatic-GFP^1-10^ and pAc5.1-SP-GFP^11^-CD4, or with. pAc5.1-Bnl^ΔHBS3^:GFP^11(7x)^ and pAc5.1-SP-GFP^1-10^-CD4. All transfections were made at 70-80% confluence in either a 6-well plate or 35 mm Glass Bottom μ-Dish. 1µg of plasmid plus 2 µL of transfection reagent were used for all single transfections; for co-tranfections 1µg of each plasmid was used plus 3µL of transfection reagent. Cells were imaged on a Olympus FV3000 inverted laser scanning confocal microscope using an oil immersion objective, UPLFLN40XO. Each cell was segmented followed by integrated density measurement and dots count on ImageJ/FIJI.

### Bnl:GFP^11(7x)^ localization analysis *in vivo*

Wing discs of L3 wandering larvae were obtained by dissection and placed live on an overhang set (Roy et al., 2014) for immediate imaging on a Olympus FV3000 inverted laser scanning confocal microscope using an oil immersion objective, UPLFLN40XO set for detection of green fluorescence. A z-stack from the basal to apical compartment of the Bnl producing cell was imaged, based on the mCherry-CAAX expression of these cells. For GFP reconstitution quantification, a SUM z projection was made and the GFP intensity was quantified in the Bnl producing cell area. The results were plotted relative to control or to wild type.

### Data analysis and statistics

All statistical analyses were performed using R (RStudio environment). Comparisons between two groups were conducted using Student’s t-tests, while experiments involving more than two groups were analyzed by one-way analysis of variance (ANOVA) followed by Tukey’s Honestly Significant Difference (HSD) post hoc test for multiple comparisons. Data visualization was performed using the ggplot2 package within the tidyverse framework. The sample size (n), definition of biological and/or technical replicates, and the specific statistical tests applied are indicated in the corresponding figure legends for each experiment. A significance threshold of p < 0.05 was adopted unless otherwise specified.

---

**Supplementary Figure 1:**
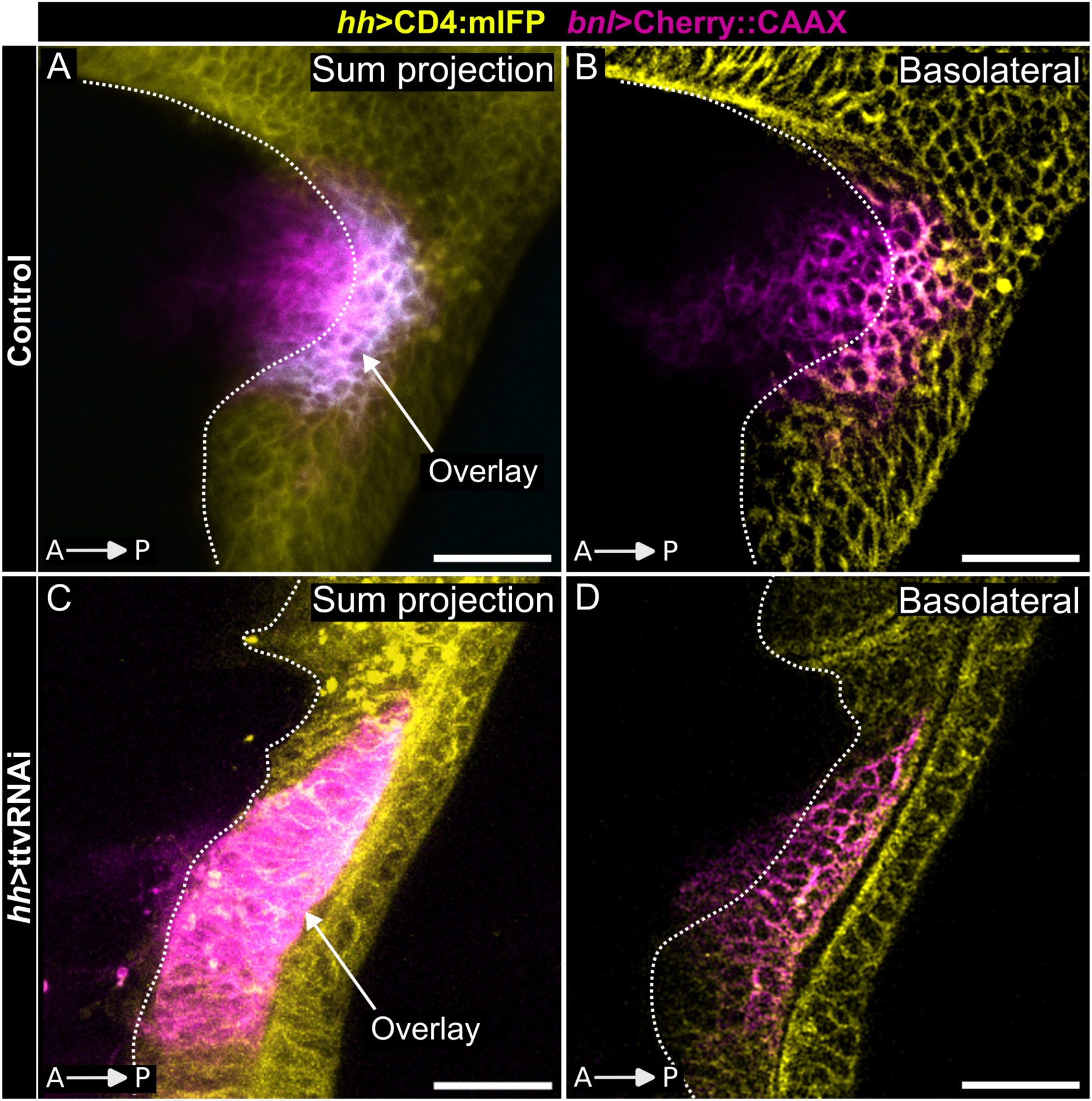
Bnl-producing domain straddles A/P compartment border. (A-), Wing disc with Bnl-producing cells expressing Cherry:CAAX (*bnl*-Gal4>UAS-Cherry:CAAX – magenta) and P compartment cells expressing mIFP (*hh*-Gal4>UAS-CD4:mIFP). (A,C) Sum projection of confocal stacks, and (B,D) basal lateral views showing Bnl-producing cells (magenta) and P compartment (yellow); overlay (white); A/P compartment border (dashed line). (A,B) controls; (C,D) disc also expressing ttvRNAi in dorsal compartment (*ap*-Gal4>UAS-ttvRNAi).

**Supplementary Figure 2:**
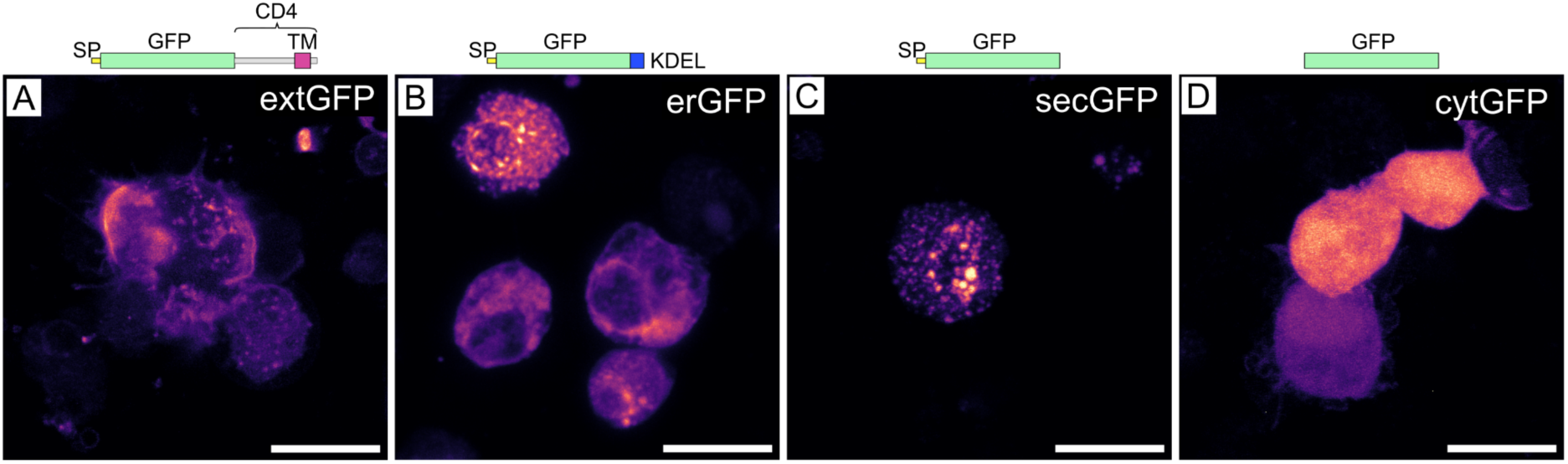
Subcellular targeting GFP in S2 cells. S2 cells expressing: (A) extGFP (GFP linked to a N-terminal signal peptide (SP) and C-terminal CD4 transmembrane domain (TM)) localizes to plasma membrane; (B) erGFP (erGFP linked to N-terminal SP and a C-terminal KDEL sequence) localizes to ER; (C) secGFP (GFP with N-terminal SP) localizes in cytoplasmic puncta; (D) cytGFP (GFP alone) localizes in cytoplasm.

**Supplementary Figure 3:**
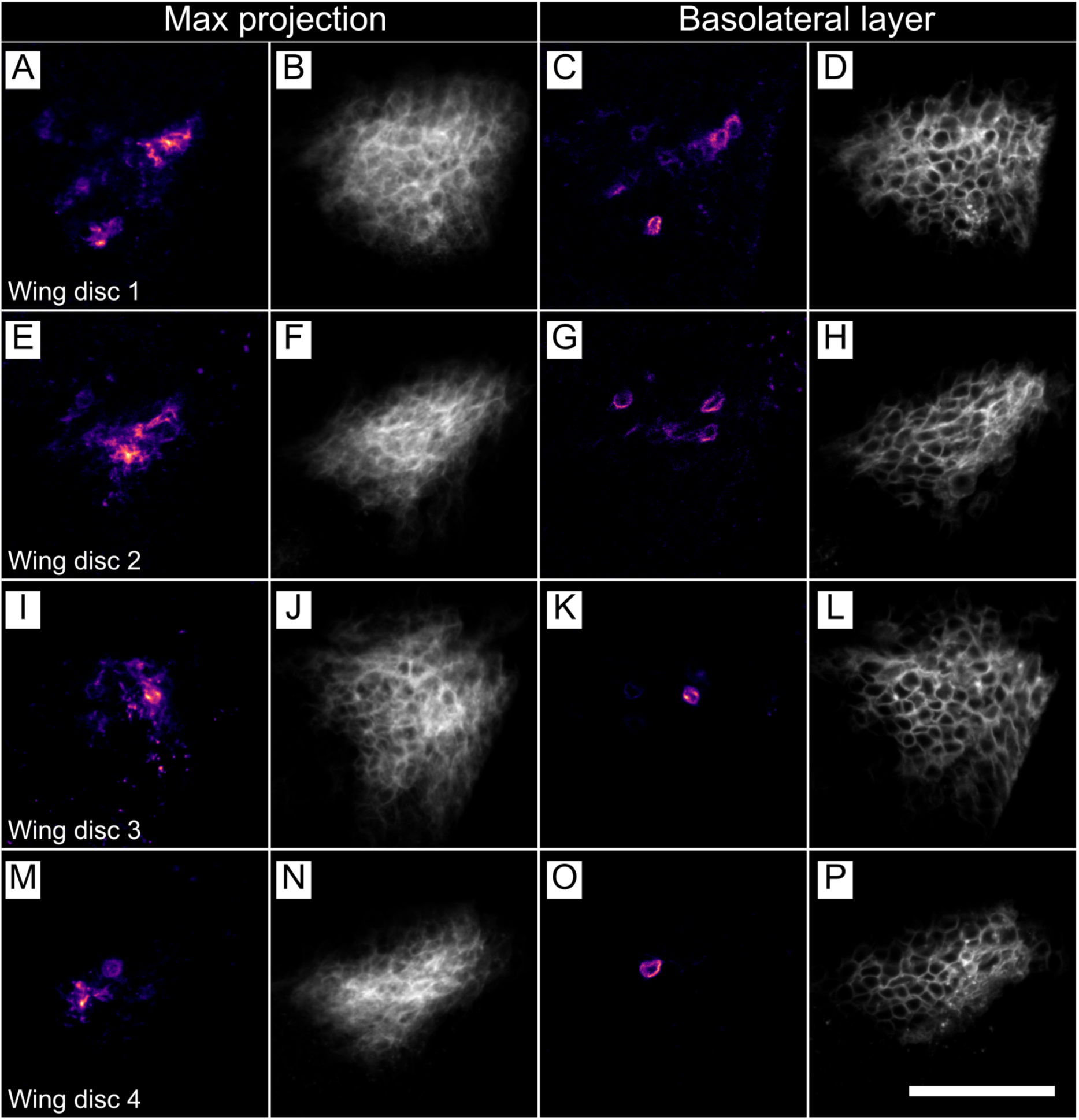
Cells with external Bnl in the Bnl-expressing domain of the wing disc. Four wing discs, genotype *ap*-Gal4/+;Bnl-LexA lexOp-mCherry:CAAX/UAS-CD4:GFP^11^, lexOp-Bnl^WT^GFP^11(7x)^. Max projections of GFP fluorescence of reconstituted Bnl:GFP (A,E,I,M) and of mCherry-marked membranes (C,G,K,O) in Bnl-expressing cells. Confocal sections with GFP fluorescence of reconstituted Bnl:GFP (C,G,K,O) and of mCherry-marked membranes (D,H,L,P) in Bnl-expressing cells. Scale bar, 40 µm.

